# Nutritional and genetic variations within foxtail millet (*Setaria italica*) landraces collected from indigenous communities across the Philippines

**DOI:** 10.1101/2023.05.22.541853

**Authors:** Nelzo C. Ereful, Jose Arnel O. Reyes, Don Emanuel M. Cardona, Roneil Christian S. Alonday, Nel Oliver B. Mateo, Huw Jones, Lesley Boyd, Evelyn F. Delfin

## Abstract

Unknown to many, the Philippines is host to a few remaining accessions of the underutilized and understudied cereal foxtail millet *(Setaria italica* (L.) P. Beauv.). We collected together accessions from different eco-geographical locations within the Philippines, along with a few accessions from Lanyu, Taiwan, to undertake a study of the nutritional value and genetic diversity within accessions of foxtail millet grown in the Philippines. All accessions were field grown in 2022, dry season (DS) at the Institute of Plant Breeding (IPB) Experiment Station, Los Baños, Laguna, Philippines. The accessions were tested for micronutrients, including Zn and Fe, nitrogen as a proxy for protein, β-carotene and a number of phenolic compounds with known nutritional potential. Of the 20 accessions tested, the accessions Bayaras and GB61438 had the highest level of Zn (107.1 mg/kg) and Fe (70.52 mg/kg), respectively, higher than levels found in traditional rice varieties. For β-carotene the highest concentration was found in the accession Balles (∼10µg/g). Twelve phenolic compounds were detected, with catechin, syringic acid, ferulic acid and kaempferol having the highest concentrations and greatest variation between accessions. To assess the genetic diversity of these local foxtail millet accessions, we sequenced a core set of eight accessions to a depth of at least 25-fold. Analysis of the population structure, using genome-wide, high-quality SNPs, showed modest diversity among the accession, with two unadmixed groups. The accessions are monophyletic relative to their earliest common ancestor, with the very light brown accessions emerging earlier than the light brown and reddish-brown varieties. Analysis of Zinc/Iron permease (ZIP) transporters within the foxtail millet reference sequence, var. Yugu1 identified 17 putative ZIP transporters. Variant calling identified SNPs primarily within 3’ and 5’ regions, and introns, indicating variation between foxtail millet accessions within regulatory gene regions rather than in structural proteins. The local foxtail millet accessions found across the Philippines, therefore, represent a potential alternative source of nutrition that would help to address the problems of the double-burden of malnutrition found in the Philippines.

## Introduction

Among the more than 100 species within the genus *Setaria*, only foxtail millet (*S. italica* (L.) P. Beauv.) became an important domesticated crop [1]. It is a self-pollinating diploid (2n = 2x = 18) with a small genome size (∼490 Mb [2]). It is easy to grow, having a short growing period, with good productivity, even under conditions of abiotic stress. It is quickly becoming a model organism in which to study beneficial traits such as C_4_ evolution, biofuel potential and the molecular mechanisms underlying abiotic and biotic resistance [3]. Foxtail is cultivated for food and fodder across Europe, China and other Asian countries. The crop is also grown in Australia, North and South America, and parts of Africa as a minor cereal [4]. Archaeobotanical evidence supports the significant contribution of foxtail millet in ancient civilization [5]. It was thought to originate in China, where domestication began, becoming a staple crop in Eurasia [1].

As a cereal crop, foxtail millet has been shown to contain comparable and often higher nutrient content (protein, dietary fibre, Ca, Zn, Fe, etc.) relative to major cereals such as wheat and rice [6; 7]. Thus, this crop could be exploited as an alternative source of nutrients, particularly in the Philippines where people suffer from widespread malnutrition, the most alarming of which is iron deficiency affecting more than half of the population [8]. A study of foxtail millet landraces from across India grain was shown to contain mean values of 12.3 % protein, 4.7 % fat, 60.6 % carbohydrates and 3.2 % ash [9]. Nutritional composition analysis of 75 elite germplasms from India, USA and China exhibited mean values of 12.63% protein, 3.52% fat, 2.07% crude fiber, 72.89% carbohydrate and 2.30% total minerals [10], close to the values reported in the study of Indian accessions [9].

In terms of genetic variation, 62 varieties collected from Asia, Europe and Africa clustered into five groups using RFLP (digested with *Eco*R V) [11]. A study using inter simple sequence repeat (ISSR) markers identified a high level of polymorphism among 81 accessions of nine *Setaria* species [12]. This study further confirmed northern China as the centre of domestication. Gene flow among accessions from different eco-geographical areas was also shown to be high. A study conducted in Nepal also showed marked diversity among 27 foxtail millet accessions [13].

A draft genome reference of foxtail millet (∼423 Mb) has been assembled using a combination of whole genome shotgun and next generation sequencing [14]. This study identified chromosome reshuffling events and rearrangements [14]. This draft genome sequence will aid the identification of functionally important variants and future breeding design. An extensive library of genomic variations has been established from the re-sequencing of 312 foxtail millet accessions. This study identified geographical adaptation, which may facilitate the genetic improvement of the crop [15].

In the Philippines, the cultivation of this heirloom crop is now restricted to the indigenous communities of (i) Batanes, a group of small islands located at the northernmost tip of the country situated between Luzon, Philippines and Taiwan, and (ii) Northern Mindanao, including Bukidnon, Misamis Oriental and Misamis Occidental [16]. A few accessions have also been found in Cagayan and Catanduanes. How and when these accessions were introduced to these different areas is still unknown.

Limited research has been undertaken on Philippine foxtail millets. It has been suggested that the Batanes accessions are similar to accessions from Orchid (Lanyu Township) [17], an island belonging to Taiwan. This was later confirmed by an investigation of the molecular differences of Asian and European landraces using RFLP, the Batanes accessions clustering with Taiwanese varieties [11]. In another study [18] foxtail millets from Batanes were classified as having round type grains. No further studies are known to have been performed on foxtail millet landraces from the Philippines.

Foxtail millet has been neglected and is unknown to many Filipinos. The dwindling cultivation of foxtail millet is largely due to the convenient availability of easy-to-cook rice, which is financially supported and distributed by the government in the Philippines. This is compounded by the lack of conservation efforts by both local and national stakeholders, foxtail millet being a non-priority and underutilized crop. Additionally, problems associated with threshing and milling further contribute to this crop’s demise, with no mechanized approach having been introduced in the indigenous communities. Today, the Ivatans, the locals of Batanes, prepare foxtail millet as a dessert dish. In Northern Mindanao it is cooked as “suman” (millet cake wrap) and “biko” (pudding) [16]. To our knowledge, no study has been undertaken to assess the genetic diversity and nutritional value of Philippine foxtail millet accessions.

There is a need to assess the potential of foxtail millet as an alternative nutrient resource. In this paper we (1) quantify the nutritional composition of several foxtail millet accessions grown at several localities in the Philippines, with a particular focus on Zn and Fe, and (2) identify the molecular variations and diversity present among eight selected local accessions using whole-genome sequencing (WGS). Additionally, as the Philippines suffers chronic problems of Zn and Fe deficiency, we analysed the genetic variations of Zinc/Iron Permease (ZIP) transporters among the varieties, as this may shed insights on the molecular mechanism of Zn/Fe uptake.

In time with the celebration of the International Year of Millet, the project was conceptualized to motivate farmers to utilize and cultivate this ancient plant. As this heirloom is under-utilized, we hope to incite curiosity among biologists, breeders and farmers, helping to preserve this crop.

## Methods

### Plant materials

Seeds of twenty foxtail millet accessions from Batanes were collected by Dr. Rodora Hontomin and Mr. Cesar Hostallero. The Batanes varieties were tentatively named after either the donor or the field site where they were collected. One accession was collected from Amulung, Cagayan, named here as Palacu, the village site of its collection. Another variety was collected from Virac, Catanduanes, Bicol Region, which is locally known as “Ukig”. Two Philippine varieties, GB61483 and GB61739 of unknown provenances, were requested from the National Plant Genetics Resources Laboratory of IPB. Six accessions from Orchid Is., Taiwan, were requested from the Germplasm Resource Information Network (GRIN), USA. These were originally collected in 1977 from Orchid Is., Lanyu Township, Taiwan. Taiwanese accessions were included to compare their agro-morphological (data not included in this paper) and nutrient content with the Batanes accessions, since previous studies showed their potential genetic relatedness [11, 17]. Seeds of two accessions from Bukidnon were collected by Dr. Raquel Salingay, Central Mindanao University. Taken all together, there were 24 accessions collected across the Philippines, with the addition of six accessions from Taiwan.

In terms of elevation, Batan Is. of Batanes is located at the northernmost tip of the Philippines and is around 77 ft (23 m) above sea level (asl); Amulung, Cagayan, 78.2 feet (23.8 meters); Bukidnon, 3,002 ft (915 meters) asl. Laguna, the experimentation site, has an estimated elevation of 387 ft (118 m) asl. All accessions were sent to IPB, Los Baños, Laguna for agro-morphological assessment, nutritional composition analysis and Whole Genome Sequencing (WGS) analysis of a core set of accessions.

### Greenhouse trial

All accessions were initially grown in two replicates under greenhouse conditions on 31 August 2021 to propagate seed for subsequent field experiment. Greenhouse trials also aimed to (i) assess their adaptability to the current site (Laguna) since these accessions were collected from different eco-geographical locations, (ii) check for any insect pest and disease infestation or susceptibility, (iii) identify variation in traits such as growth habit, spike colour, and flowering time. Ten seeds from each accession were directly seeded into a soil mix composed of sand: silt: clay proportions of 32% – 48% – 20%, and loam as textural grade and water holding capacity (WHC) of 67%. Fungicide was applied a week prior to seeding. Seedling thinning was performed to retain 3–5 stalks. Plants were watered with approximately one litre of water once a week, as previously described [19].

Of the 30 accessions grown, five did not germinate (non-viable) at the current site. These include three Taiwanese and two Bukidnon accessions. Additionally, five accessions were discarded due to their agro-morphological similarities to other accessions (data on agro-morphological, quantitative and qualitative, will be a subject of a separate paper).

### Dry (DS) and Wet Season (WS) field experiments

Seeds of the remaining 20 accessions were grown in the field, in three block–replicates. These lines were pre-germinated in seedling trays on 14 January 2022 and transplanted on 03 February 2022 for DS field trials. The experiment was performed at E3–West of the IPB Experiment station 14°08’24.8“N, 121°15’32.3”E. The previous crop cultivated on this site during the 2021 WS was eggplant. Sugarcane was also grown in two previous season–years (2019, 2020). We implemented incomplete block design with three blocks, with accessions randomly arranged as designed by the block design application [20] in Rstudio. The field was laid out with plot dimensions of 11m × 3m, with 0.5 m in between rows and 0.25 m in between hills in each row (i.e., 50 cm × 25 cm plant spacing). Each block was separated by 1.5 m alleys. Furrow (∼15 cm in depth) and ridge sowing were implemented in DS and WS trials, respectively. Three seedlings were planted per hill with twelve hills in each row. Seedling thinning was performed, if necessary, to retain three seedlings per hill. Spikes from 10 hills in each row were collected for nutrient composition assessment. Border rows were also included in the design, but nutrient composition was not analysed from samples collected from border rows.

As practiced by the locals and to avoid bias in micro-nutrient accumulation, no fertilizer was added throughout the experiment. Pest, weed, and disease infestations were monitored regularly, and control measures were applied if needed. However, there was visually negligible incidence of disease infestation in our field trials. This is consistent with the literature, in which foxtail millet accessions are reported to be both biotic- and abiotic-stress resistant (reviewed in [21]). Weeding was performed as needed. Unless precipitation occurred, watering was done once a week for the DS as previously suggested for foxtail [19]. No watering was performed during the WS.

For the WS, seeds were sown on 8 July 2022 on seedling trays and were transplanted 20 DAS. However, the foxtail millet accessions were observed not to thrive well under wet conditions presumably due to frequent precipitation and enormous cloud cover. This is not surprising as foxtail is known to thrive under arid and semi-arid conditions and was said to be a summer crop [22]. No data on nutrient composition was, therefore, obtained for the 2022 WS.

### Soil sampling

Soil samples were collected by composite sampling before transplanting. Individual soil cores were collected using a soil auger from five points in X shape at 0–25 cm depth in each block and were mixed (three reps, in total). The representative samples were then bulked, air-dried, sieved through 2-mm mesh, and prepared for analyses at the Plant Physiology Laboratory, IPB and Analytical Services Laboratory, Department of Soil Science, Agricultural Systems Institute (ASL-DSS-ASI), College of Agriculture and Food Science, UPLB. The following were analysed: pH, organic matter content, total nitrogen (N), available phosphorus (P), potassium (K), zinc (Zn), and iron (Fe) based on procedures previously described (see Table 1 for references).

**Table 1.**
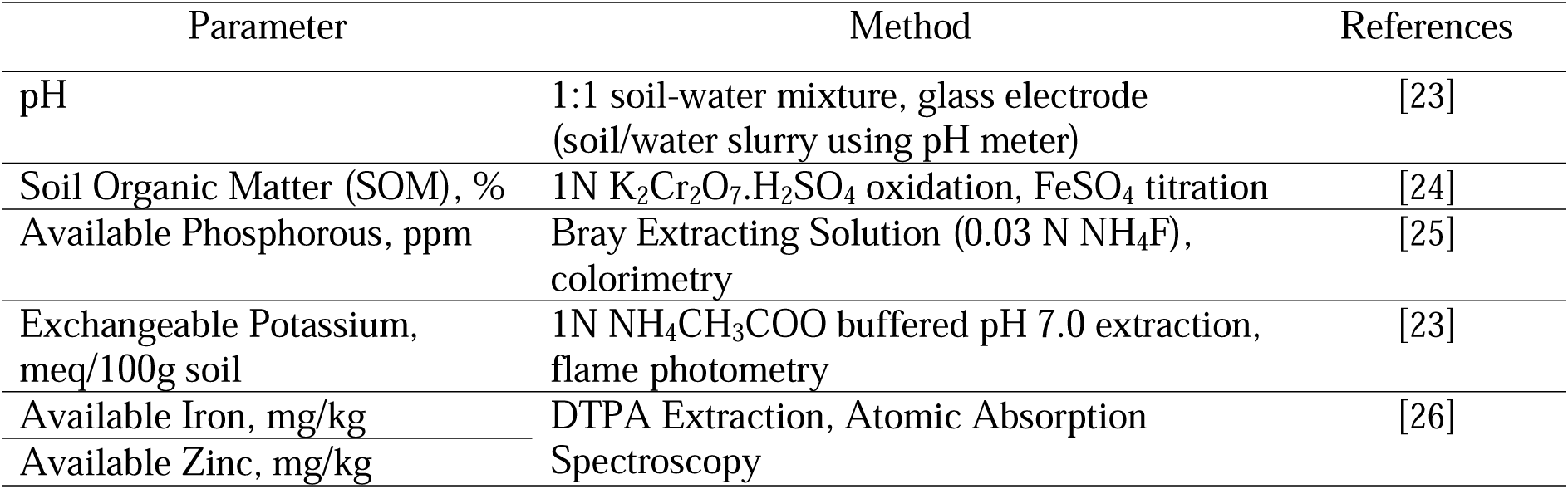
Methods of analyses and references used for determining soil chemical properties.

### Nutrient composition analysis

Grain samples were milled using an electronic grinder (RainPhil^®^) set at a fixed rotational speed for 30 seconds. Foxtail millet flour was separated from the bran, implementing manual winnowing to initially remove the hulls, then mesh-sieving with No. 80 mesh testing sieve (180 µm). Flour samples were analyzed for nutrient composition including Zn, Fe, Mg, Ca, Cu, N (proxy for crude protein) and β-carotene across the three biological reps collected from the 2022 DS field trial, implementing three technical replicates (3 bio reps × 3 tech reps = 9 reps for each element) at the Analytical Services Lab (ASL), IPB. Each biological rep represents each block– replicate of each variety collected from the field.

Nitrogen content of the foxtail millet samples was determined following the method of [27]. For the quantification of metals (Zn, Fe, Mg, Cu and Ca), one gram of each of the samples was ashed and digested with 25 mL 37 % hydrochloric acid with 5 drops of 69 % nitric acid until 10 mL concentrate was obtained. The concentrate was diluted to 50 mL with ultra-pure water. Metal ion concentrations in the dissolved ash solution were quantitatively determined by Atomic Absorption Spectroscopy (AAS) (Model: AA-7000, Shimadzu, Japan).

The phenolic compounds were analysed at the National Institute of Agricultural Botany (NIAB), Cambridge following standard procedure as previously described [28]. Phenolic compounds were assessed using HPLC. Flour samples of approx. 2.5g were extracted into 50ml of ethanol-acetic acid (10% 1M acetic acid v/v) under reflux conditions for 2h. Extracts were stored at −20°C until analysed. The extracts were prepared for chromatography by centrifugation for 2 min at 13000 rpm, then filtrated through a 0.2μm filter. The compounds were separated using the Dionex Ultimate 3000 HPLC system. A 150mm × 4.6mm × 5μm × 100Å Kinetix C18 column was used, with a gradient 0.1% formic acid /acetonitrile mobile phase running at 0.2ml/minute (gradient of 0.95: 0.05 for 2min, then 0.72; 0.28 for 18min, 0.00; 0.10 for 28min, and then held until 45min) were used. The column effluent was monitored with a PDA detector between 200 and 600nm, with data recorded at 254nm, 280nm, 340nm and 520nm. Samples were replicated, with three biological reps and two technical reps each.

Determination of β-carotene levels was adopted from [29]. One gram of dried flour sample was added to chilled 50-mL acetone. The mixture was shaken using a vortex mixer at high speed and left to stand for 15 mins, with occasional shaking. The solution was filtered using Whatman filter paper No. 42. with the receiver submerged in an ice bath. The addition of acetone was repeated twice and the supernatants pooled. The absorbance of the extract was read at 449 nm using UV-vis spectrophotometer. β-carotene levels were determined against a standard. β-carotene was analysed in three biological reps, with three technical replicates run for each sample (3 bio reps × 3 tech reps = 9 reps).

### Statistical analysis

R (v. 4.1.0 [30]) was used to analyse datasets. For micronutrient analysis (e.g., Zn, Fe) basic statistical tests, such as means and standard deviations, were calculated using the ‘aggregate’ function. Two-way Analysis of Variance (ANOVA) was fitted in our samples using ’aov’ function to infer significance among accessions. Tukey’s HSD (Honest Significant Difference) was also performed to compare each group in a pairwise fashion.

Histograms were generated to view distribution of datasets. Residual analysis was also performed by calculating and creating plots on Residuals vs Fitted plot, Normal QQ plot, Scale– Location, Residuals vs Leverage. Datasets were also tested using Shapiro–Wilk normality test and Bartlett test of homogeneity of variances. As described previously, we ensure our statistical analyses are robustified by generating biological and technical replicates. Figures were created using ggplot2 package [31].

Correlations between trace minerals, N, and β-carotene were calculated using Pearson rank correlation in rcorr function found in Hmisc package. Likewise, correlations were also calculated between phenolic compounds.

### DNA extraction

For DNA extraction, flag leaf samples were taken at 35 days after transplanting (DAT). Until usage, samples were kept at -80°C. The modified CTAB protocol [32], was employed to separate genomic DNA from the foxtail leaf samples. Frozen plant tissues were ground up using a frozen mortar and pestle. To remove polysaccharides and other impurities, samples were treated with a sorbitol pre-wash buffer (100 mM Tris-HCl, 0.35 mM sorbitol, 5 mM EDTA, 1% (w/v) PVP, 1% (v/v) 2-mercaptoethanol) before extracting DNA with warm CTAB buffer (100 mM Tris-HCl, 3 mM NaCl, 3% CTAB, 20 EDTA, 1% (w/v) PVP, 1% (v/v) 2-mercaptoethanol). The mixtures were incubated at 65°C for 1 hour, with inversions every 10 minutes. Chloroform:isoamyl (24:1) was added, and the mixture was centrifuged. Aqueous phases were collected, and DNA was precipitated with isopropanol. DNA pellets were washed with 70% ethanol and resuspended in 50 μL of TE buffer containing 0.1 mg/mL RNase A. Quantitative and qualitative analyses were done on the extracted gDNA to determine its purity using Nanodrop™ and agarose gel electroporation. The samples were stored at –80°C until shipped.

### Whole Genome Sequencing

Genomic DNA was sequenced by Novogene, Singapore using the Illumina platform NovaSeq PE150. Briefly, gDNA was randomly sheared into short fragments, end repaired, A-tailed and ligated with Illumina adapters. The fragments were PCR-amplified, size selected and purified. Reads were checked for adapters, and low-quality reads (at Q > 20) by Novogene. No further pre-processing procedure were required as demonstrated using FASTQC [33] since there are no adapters and low-quality reads. The genome sequence of the foxtail reference sequence (*Setaria italica* cv. Yugu1 v2.0.54) was downloaded from Ensembl Plants [34] and indexed using BWA [35]. A dictionary for the reference genome was created using PICARD tool (option: CreateSequenceDictionary). Reads were mapped against the indexed genome reference using BWA (option: mem), implementing default parameters. Outputs were piped to bam and sorted using SAMtools [36]. PICARD was used to assign reads to a single Read Group (option: AddOrReplaceReadGroup) after which PCR duplicates were flagged using MarkDuplicates option. BAM files were indexed using PICARD (option: BuildBamIndex)

Variants were identified using the Broad Institute Genome Analysis Tool Kit (GATK v4.1.9.0) [37]. Best Practices for germline SNP/Indel discovery. GATK (option: Haplotypecaller) was run for each foxtail millet accession dataset, using its default parameters, to call variants between the reference genome sequence and each sequenced accession, in both Variant Call Format (VCF) and Genomic VCF (option: --ERC GVCF; multiple samples) modes. For individual samples, we used VCFtools v0.1.15 [38] to filter the variant calls with the following parameters: a Minor Allele Frequency (MAF) of 0.05 to 0.1 (the number of output did not change within the range); a minimum mean depth for a site, 7; minimum quality score 30; and, a minimum missing data of 0.9 (--maf 0.05 to 0.1 --min-meanDP 7 --minQ 30 --max-missing 0.9). Note that Indels were not removed at this stage.

Using GVCF mode for multiple samples, all accessions were merged using GATK CombineGVCFs command. Joint genotyping was performed using GATK GenotypeGVCFs. Indels (option: --remove-indels) and monomorphic SNPs were removed (option: --maf 0.001) sequentially using VCFtools. Sites were filtered using the same tool with the following options: --min-meanDP 7 - -max-missing 0.5 --mac 3 --minQ 30. This enabled us to kept variants that have been successfully genotyped in 50% of individuals, a mean depth value of 7, a minimum quality score of 30, and a minor allele count of 3. Variant Effect Predictions was performed using SNPEff 5.1d [39] and Setaria_italica as database from Ensembl.

Structure analysis. FastStructure [40] was used to infer population structure within the foxtail millet accessions using the same SNPs, with simple (at K = 2 to 5) prior.

Phylogenetic tree. Using PLINK2 the VCF file of the whole genome sequence was converted into PLINK binary format (bed, bim and fam). A phylogenetic tree was created using VCF kit [41] using the genome-wide SNPs identified across the foxtail millet accessions implementing Neighbor– Joining algorithm.

Principal Component Analysis (PCA). Using the VCF file a PCA plot was created using PLINK2. Linkage pruning and removal of rare variants was first performed. The plot was generated using the libraries tidyverse [42] and ggplot2 in R.

Phylogenetic analysis. Fasta files of both gene and translated amino acid ZIP transporter sequences of foxtail millet, rice, and *A. thaliana* were all downloaded from Ensembl Plants. Multiple sequence alignment (MSA) was performed using Clustal Omega [43] with output in Newick format. Phylogenetic tree was created using VCF-kit [41]. All scripts are run in R.

## Results

Foxtail millet accessions were collected from several eco-geographical sites across the Philippines (Table 2). These include accessions collected from Batanes (20 accessions), Bukidnon (2 accessions), Cagayan (1 accession) and Catanduanes (1 accession) (Fig. 1). Two local varieties of unknown provenances and 6 Taiwanese accessions were also included in the study (Table 2). From this collection, seeds from 20 accessions were subsequently multiplied during the 2022 DS field trial (Table 2; Fig. 2). Milled grain was analysed in these 20 accessions for elemental micronutrients, N as a proxy for crude protein content, β-carotene and a range of phenolic compounds. Whole genome sequencing (WGS) was undertaken on eight of the accessions to assess diversity.

**Table 2.**
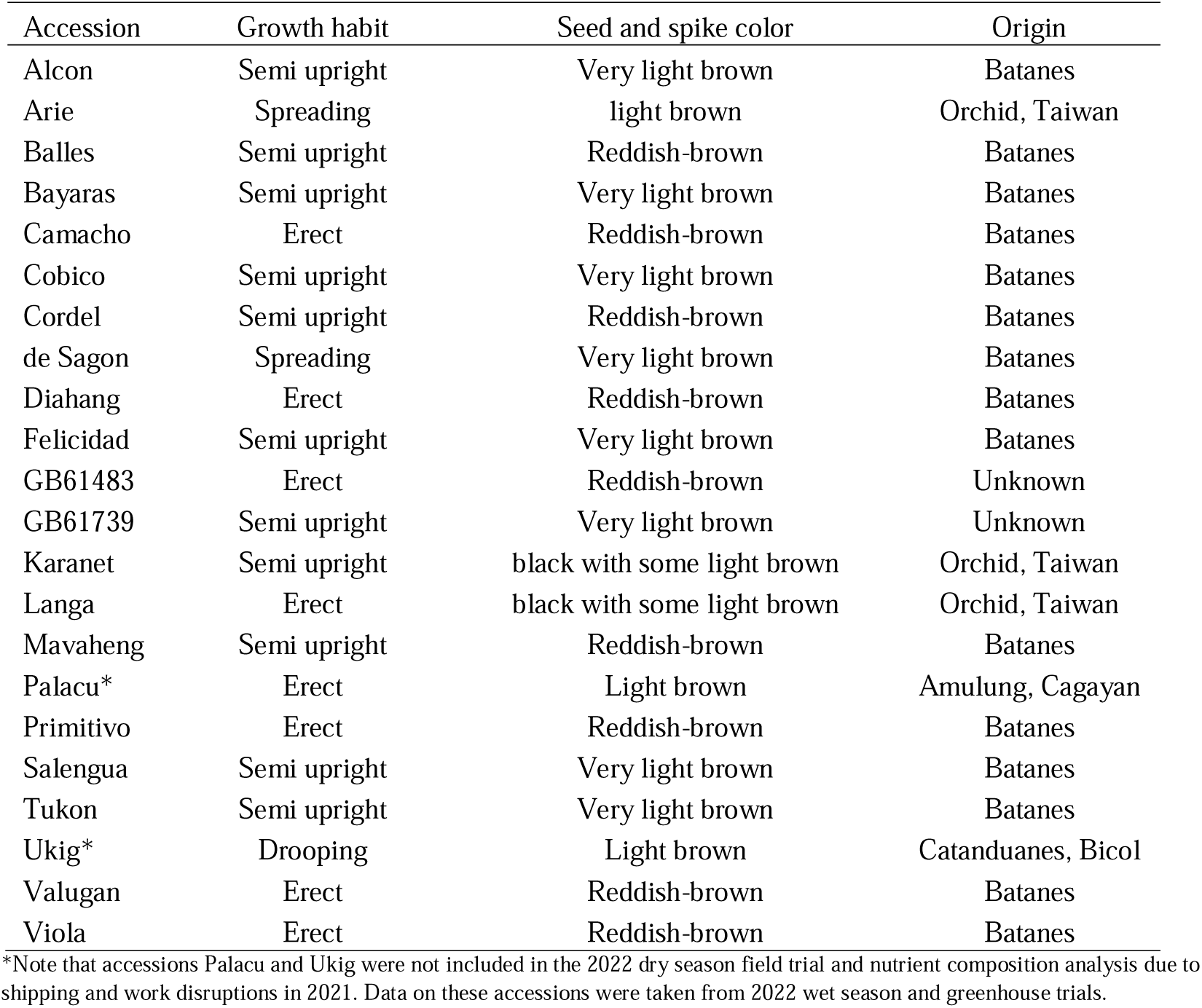
Foxtail millet accessions planted in the 2022 DS field trials: visual characteristics and origin.

**Fig. 1.**
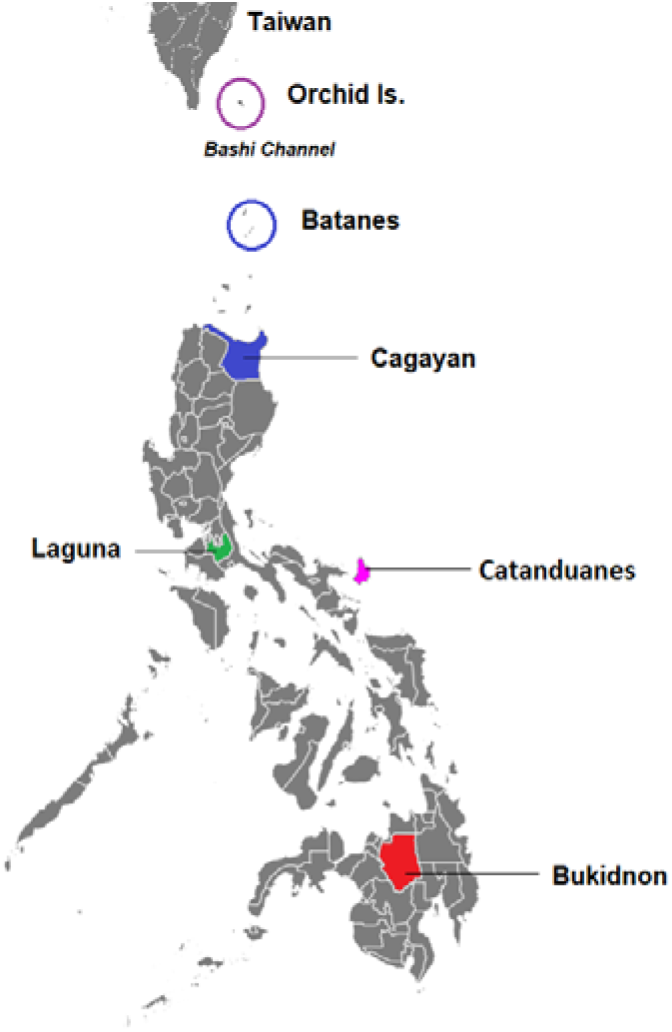
Eco-geographical locations of the foxtail millet accessions collected in this study: (i) Batanes (circled blue), located at the northernmost tip of the Philippines; (ii) Cagayan (filled blue); (iii) Bukidnon, highlighted in red, and (iv) Orchid Is. (circled purple). Also shown is Laguna (green), the site of greenhouse and field experimentation trials. Accessions from Orchid Is. were shipped from USDA and were originally collected by the Germplasm Resource Information Network (GRIN). Map taken from https://simplemaps.com.

**Fig. 2.**
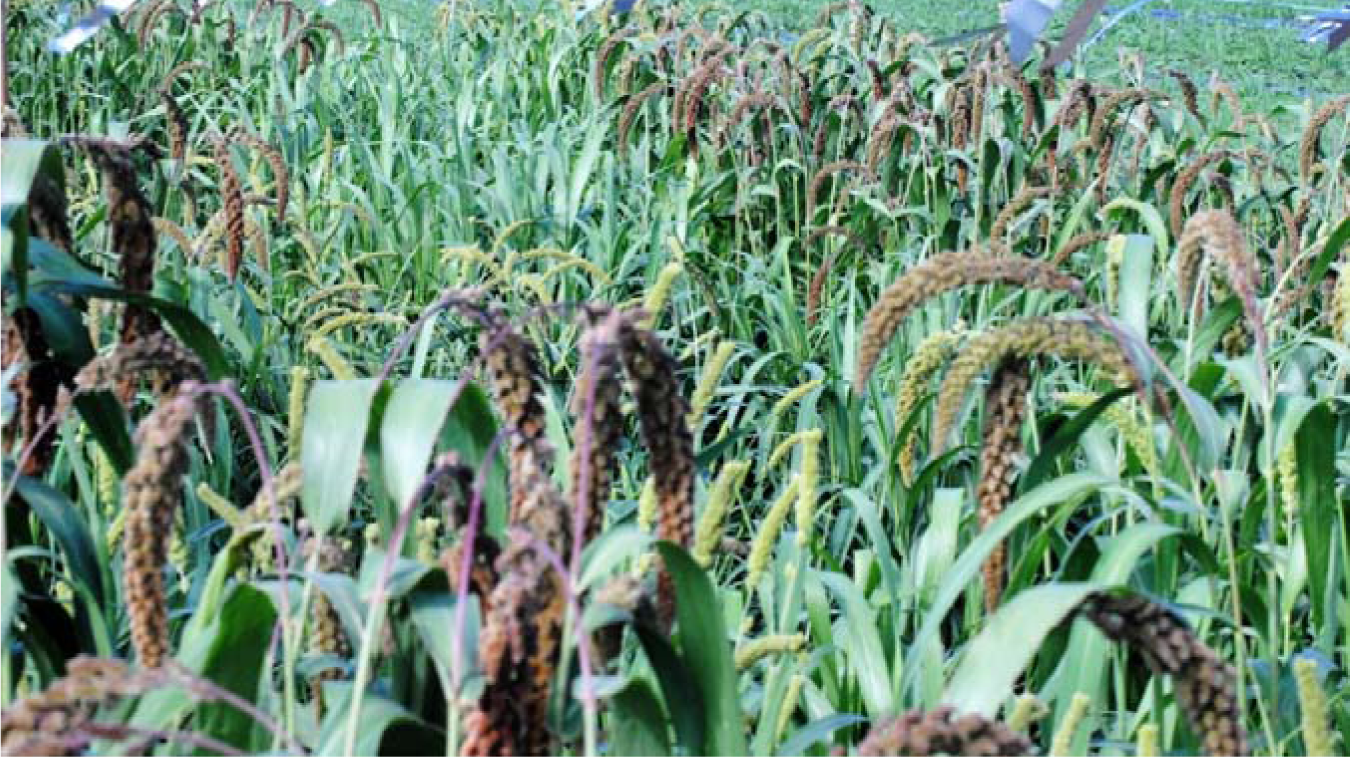
Philippine and Taiwanese foxtail millet accessions planted at the field experimentation site of the Institute of Plant Breeding (IPB) during the 2022 DS.

## Field Experiment

### Soil analysis of the field site

Analysis of the soil samples collected from the experimentation site (E3–West, IPB, UPLB) showed a composition of 31% sand, 42% silt, and 27% clay; a clay–loam textural grade; and a water holding capacity (WHC) of 66%. Specific soil analysis of the three blocks showed that the site is moderately acidic, with an average pH of 6.04 (Table 3). Organic matter (OM) content was an average of 2.01%, which is classified as a medium content scale [44]. Total N and available N showed an average of 2006.7 kg N/ha and 60.2 kg N/ha, respectively. Available P concentration was high at 15.62 ppm; K levels were of medium concentration at 0.44 meq/100g soil. The field site had a medium concentration of Zn and high levels of Fe, with an average 4.78 mg/kg and 104.10 mg/kg, respectively, based on the classification by [44].

**Table 3.** Soil analysis of E3, IPB, the site of the field experiment for the 2022 DS.

| Block | pH<br>(1:1, water) | OM<br>(%) | Total N<br>kg N/ha | Available N<br>kg N/ha | Avail P<br>Bray P2 (ppm) | K<br>meq/100g | Zn<br>(mg/ kg) | Fe<br>(mg/ kg) |
| --- | --- | --- | --- | --- | --- | --- | --- | --- |
| 1 | 6.12 | 2.07 | 2070 | 62.1 | 16.18 | 0.387 | 3.99 | 116.18 |
| 2 | 5.97 | 1.94 | 1940 | 58.2 | 13.27 | 0.412 | 5.14 | 104.06 |
| 3 | 6.04 | 2.01 | 2010 | 60.3 | 17.42 | 0.511 | 5.22 | 92.05 |
| Ave | 6.04 | 2.01 | 2006.67 | 60.20 | 15.62 | 0.44 | 4.78 | 104.10 |

Note: OM, organic matter; N, nitrogen; P, phosphorus; K, potassium; Zn, zinc; Fe, iron

### Nutritional composition analysis of foxtail millet accessions

Milled grain from 20 foxtail millet accessions were analysed for N, as a proxy for crude protein content, the micronutrients Fe, Zn, Mg, Ca and Cu, β-carotene and a range of phenolic compounds. Grain of the 20 accessions used for these assessments was obtained from the plants grown at the field experimentation site during the DS. Eight of these accessions were analysed for WGS.

### Ash and Nitrogen analyses

Ash content of the accessions ranged from 0.59% (GB61739) to 1.07% (Camacho), with an overall average across accessions of 0.84% (0.84 g/100g) (see Suppl. Table 1). Analysis of the N content, as a proxy to protein content, showed significant differences (ANOVA, *P* < 0.001) among the accessions, with a mean value of 1.67 g/100g (or 16.7 g/kg) (Fig. 3; Suppl. Table 2). The accessions de Sagon and Camacho exhibited the lowest (0.90 g/100g) and highest (2.14 g/100g) concentrations of N, respectively, with 12 of the accessions have high N levels, similar to Camacho (Fig. 3). Based on a nitrogen-to-protein conversion factor (NPCF) of 6.25, the crude protein content would range from 5.63 to 13.38 g/100g (or 5.63 to 13.38%).

**Fig. 3.**
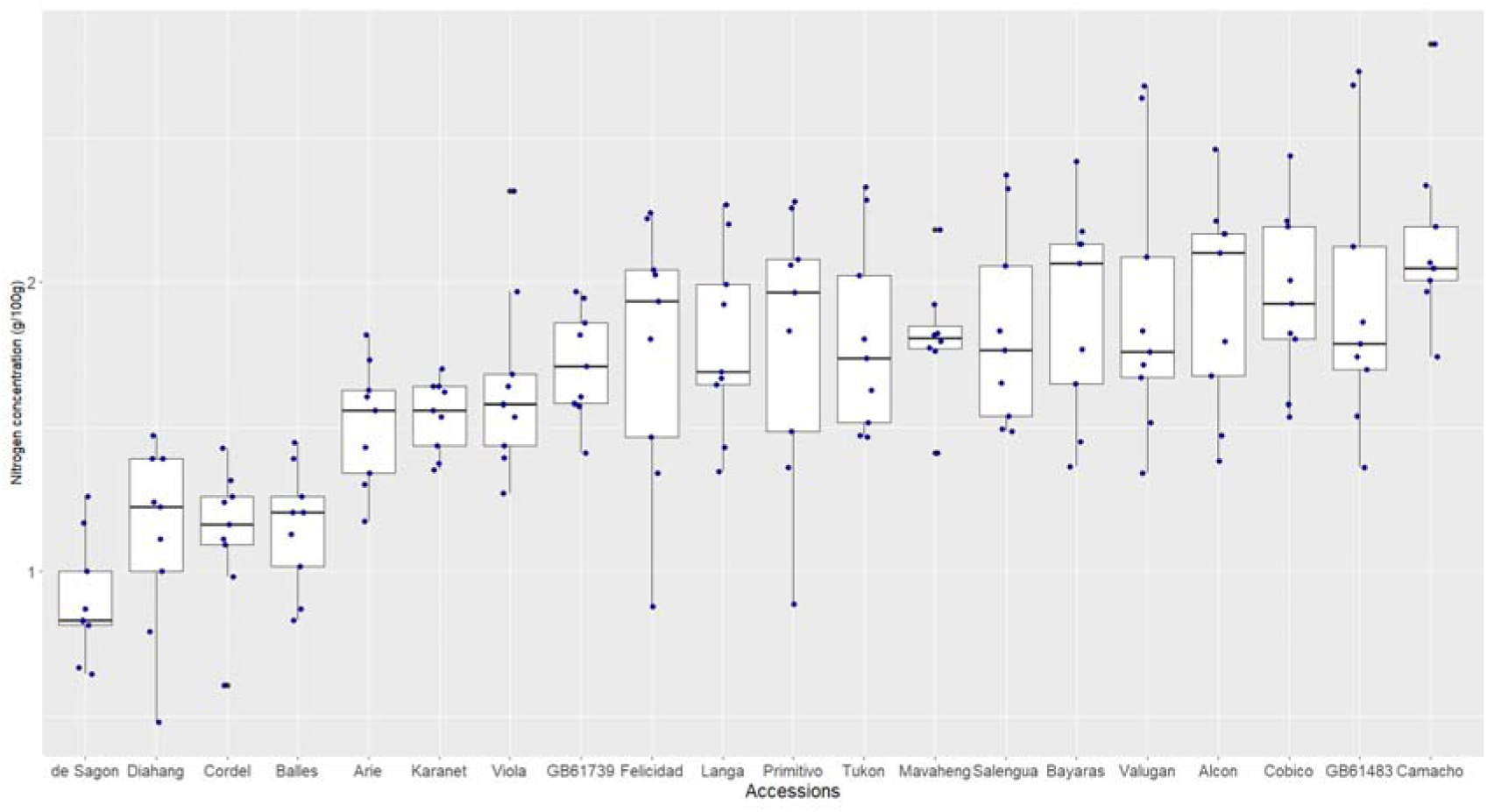
Variation in nitrogen content in 20 foxtail millet accessions.

### Micronutrient content (Zn and Fe)

Focusing on minerals essential for human health, we analysed Zn and Fe levels in the 20 accessions. Significant variation in Zn concentrations was observed (ANOVA, *P* < 0.001) with overall mean of 52.60 mg/kg across all accessions, with average values ranging from 34.63 mg/kg (GB61739) to 107.05 mg/kg (Bayaras) (Fig. 4; Suppl. Table 3 for raw data). Compared to the three Taiwanese accessions (Arie, Karanet and Langa), the Philippine lines Tukon, Felicidad, Valugan, and Bayaras were found to contain significantly higher levels Zn (Tukey’s HSD; Fig. 4; Suppl. Table 4).

**Fig. 4.**
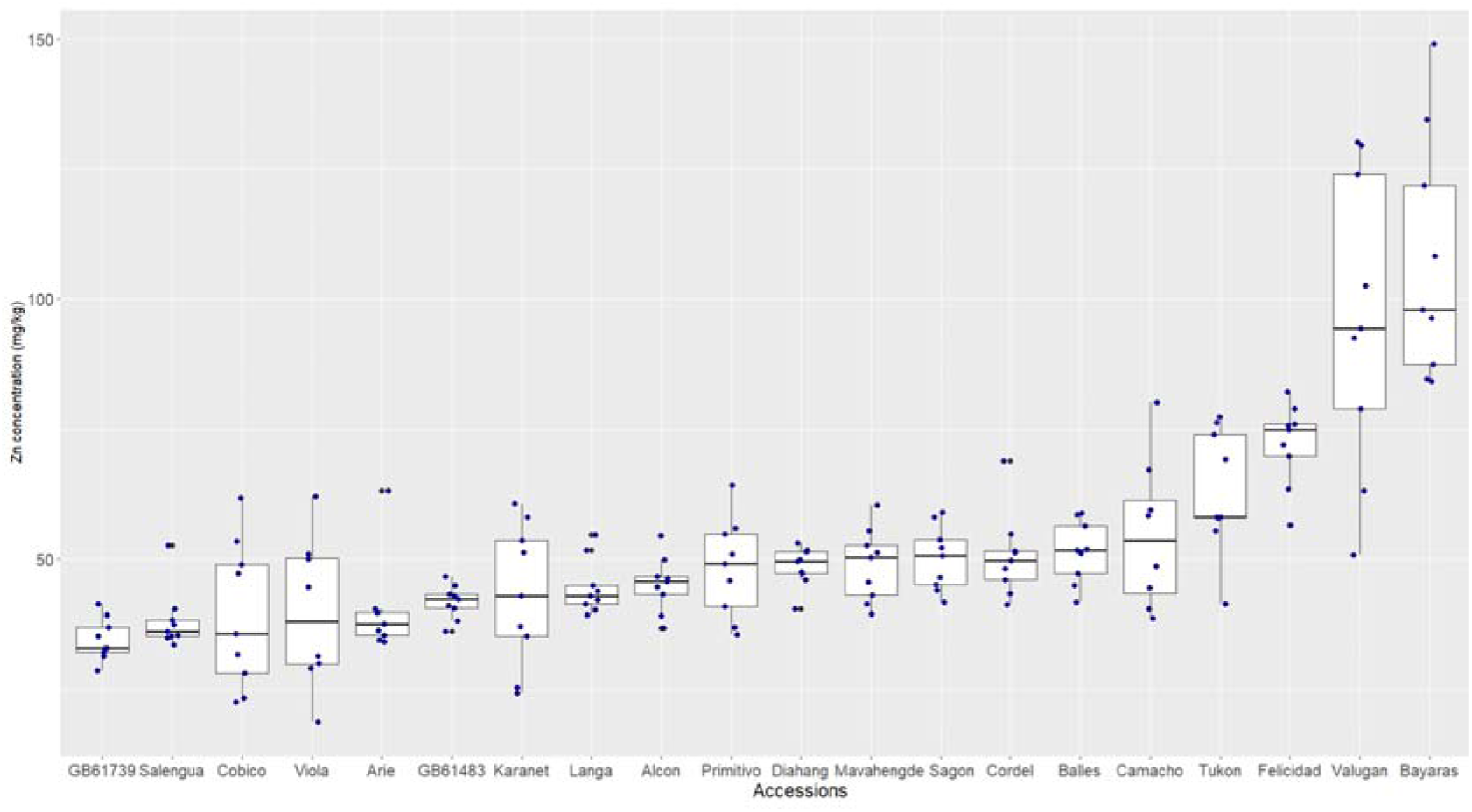
Variation in Zn (mg/kg) content in 20 foxtail millet accessions.

Similarly, significant variation in Fe concentrations was observed across the accessions (ANOVA, *P* < 0.001). The mean concentration of Fe was 45.66 mg/kg. The accessions Felicidad and GB61483 were found to contain the lowest and highest mean concentrations of Fe, with 30.04 and 70.52 mg/kg, respectively (Fig. 5; Suppl. Table 5).

**Fig. 5.**
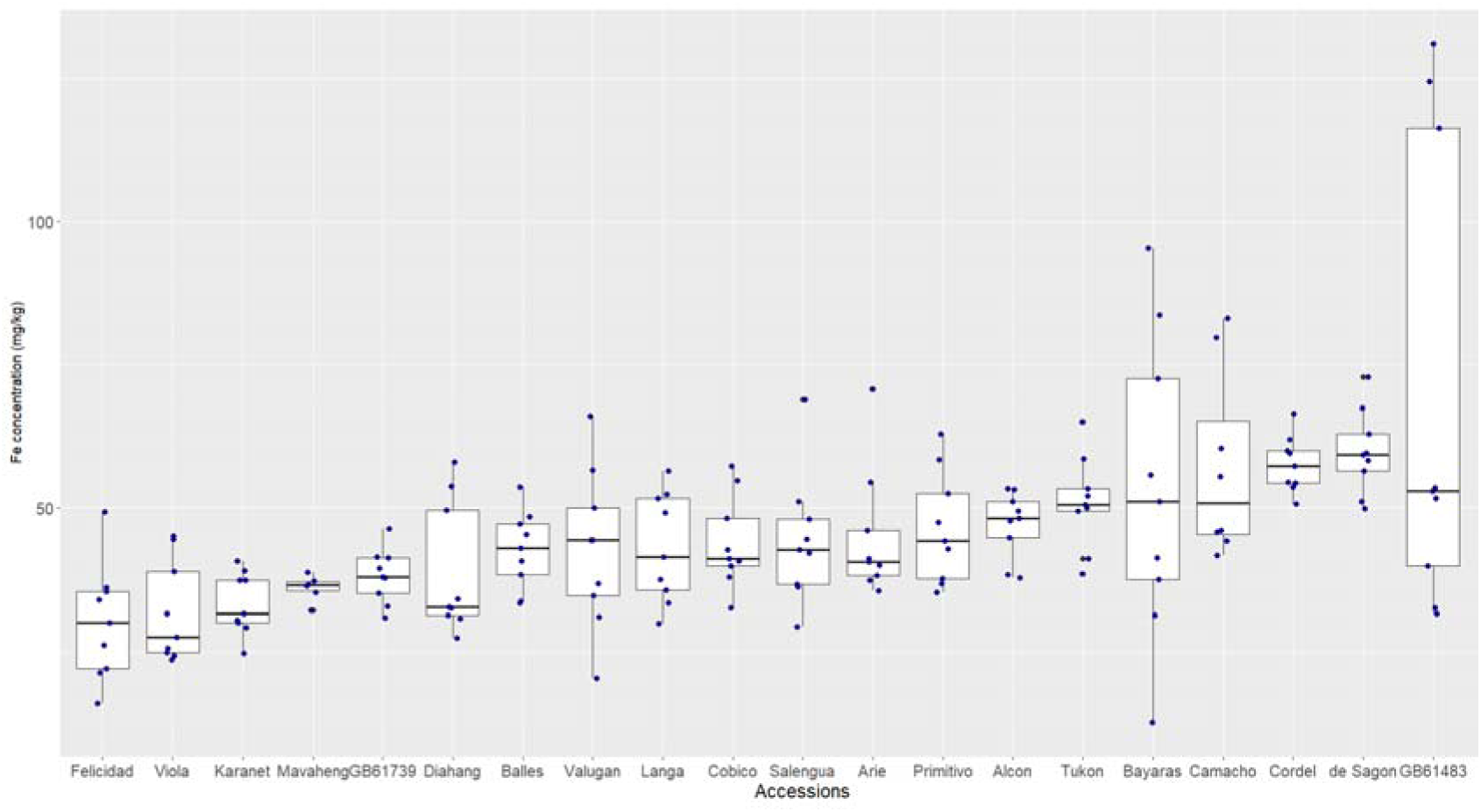
Variation in Fe (mg/kg) content in 20 foxtail millet accessions.

### Other micronutrient concentrations (Mg, Ca and Cu)

Analysis of Mg, Ca and Cu concentrations showed significant variations between the foxtail millet accessions (*P*<0.001). Mg had a mean value of 282.84 mg/kg (Table 4). Primitivo was found to contain the lowest concentration of Mg with 154.29 mg/kg and Diahang, the highest, at 595.57 mg/kg (Suppl. Table 6). The accessions demonstrated a mean Ca concentration of 564.14 mg/kg, with values ranging from 229.04 mg/kg in accession GB61483 to 1504.59 mg/kg in Bayaras (Table 4; Suppl. Table 7). Cu concentration had a mean value of 7.41 mg/kg. The accessions Valugan and Camacho showed the highest concentrations of this trace mineral, with 10.18 and 9.93 mg/kg, respectively. Accession GB61739 had the lowest concentration, with 3.73 mg/kg (Table 4; Suppl. Table 8).

**Table 4.** Average Mg, Ca, and Cu (in mg/kg) contents in 20 foxtail millet accessions*.

| Accessions | Mg | Ca | Cu |
| --- | --- | --- | --- |
| Diahang | 595.57<sup>a</sup> | 481.24<sup>d</sup> | 9.56<sup>bc</sup> |
| Cordel | 500.06<sup>b</sup> | 438.34<sup>de</sup> | 8.79<sup>d</sup> |
| de Sagon | 461.79<sup>b</sup> | 458.94<sup>de</sup> | 8.84<sup>d</sup> |
| Bayaras | 362.40<sup>c</sup> | 1504.59<sup>a</sup> | 8.99<sup>cd</sup> |
| Balles | 317.08<sup>d</sup> | 450.22<sup>de</sup> | 7.03<sup>f</sup> |
| Felicidad | 287.21<sup>de</sup> | 1147.97<sup>b</sup> | 8.00<sup>e</sup> |
| Valugan | 285.76<sup>d-f</sup> | 1399.82<sup>a</sup> | 10.18<sup>a</sup> |
| Arie | 275.24<sup>e-g</sup> | 268.71<sup>g-i</sup> | 5.91<sup>h</sup> |
| Salengua | 257.52<sup>e-h</sup> | 250.16<sup>hi</sup> | 7.71<sup>e</sup> |
| Viola | 251.21<sup>e-i</sup> | 1093.47<sup>b</sup> | 6.26<sup>gh</sup> |
| Tukon | 245.91<sup>f-j</sup> | 304.90<sup>g-i</sup> | 7.91<sup>e</sup> |
| Langa | 236.98<sup>g-j</sup> | 354.29<sup>e-h</sup> | 3.79<sup>i</sup> |
| Mavaheng | 223.41<sup>h-j</sup> | 289.36<sup>g-i</sup> | 6.22<sup>gh</sup> |
| Camacho | 213.59<sup>i-k</sup> | 315.54<sup>f-i</sup> | 9.93<sup>ab</sup> |
| GB61739 | 211.54<sup>i-k</sup> | 302.68<sup>g-i</sup> | 3.73<sup>i</sup> |
| GB61483 | 210.60<sup>jk</sup> | 229.04<sup>i</sup> | 8.62<sup>d</sup> |
| Alcon | 210.39<sup>jk</sup> | 426.29<sup>d-f</sup> | 6.41<sup>g</sup> |
| Cobico | 178.45<sup>kl</sup> | 312.68<sup>g-i</sup> | 9.54<sup>bc</sup> |
| Karanet | 177.89<sup>kl</sup> | 878.69<sup>c</sup> | 5.85<sup>h</sup> |
| Primitivo | 154.29<sup>l</sup> | 375.90<sup>d-g</sup> | 6.98<sup>f</sup> |
| Average | 282.84 | 564.14 | 7.51 |

\*Means in each column followed by the same letter(s) are not significantly different at 5% level using Least Significant Difference (LSD) test for multiple comparisons. Table sorted based on the values of Mg.

**Table. 4.** Summary of variant calls (SNPs and Indels) in each accession after filtering.

| Varieties | Total no. of variants | No. of SNPs | No. of Insertions | No. of Deletions | Variant Rate (for every 1 kb) |
| --- | --- | --- | --- | --- | --- |
| Palacu | 920,357 | 842,834 | 35,593 | 41,930 | 2.27 |
| Mavaheng | 863,841 | 790,133 | 33,685 | 40,023 | 2.14 |
| Felicidad | 903,157 | 826,214 | 35,545 | 41,398 | 2.23 |
| Bayaras | 833,621 | 833,621 | 32,385 | 38,198 | 2.06 |
| GB61739 | 826,818 | 787,481 | 33,601 | 39,742 | 2.13 |
| GB61438 | 860,824 | 756,766 | 32,254 | 37,798 | 2.04 |
| De Sagon | 943,601 | 863,265 | 37,020 | 43,316 | 2.33 |
| Ukig | 968,517 | 886,292 | 37,631 | 44,594 | 2.39 |

### β-carotene content

All 20 accessions of foxtail millet were found to contain β-carotene, with an overall mean of 5.39 µg/g. The accession Arie had the lowest β-carotene concentration, with 3.44 µg/g (Fig. 6; Suppl. Table 9). On the other hand, Balles and Diahang showed the highest levels of β-carotene concentration, with 9.69 and 9.12 µg/g (not statistically significant using Tukey’s HSD; Suppl. Table 10), respectively.

**Fig. 6.**
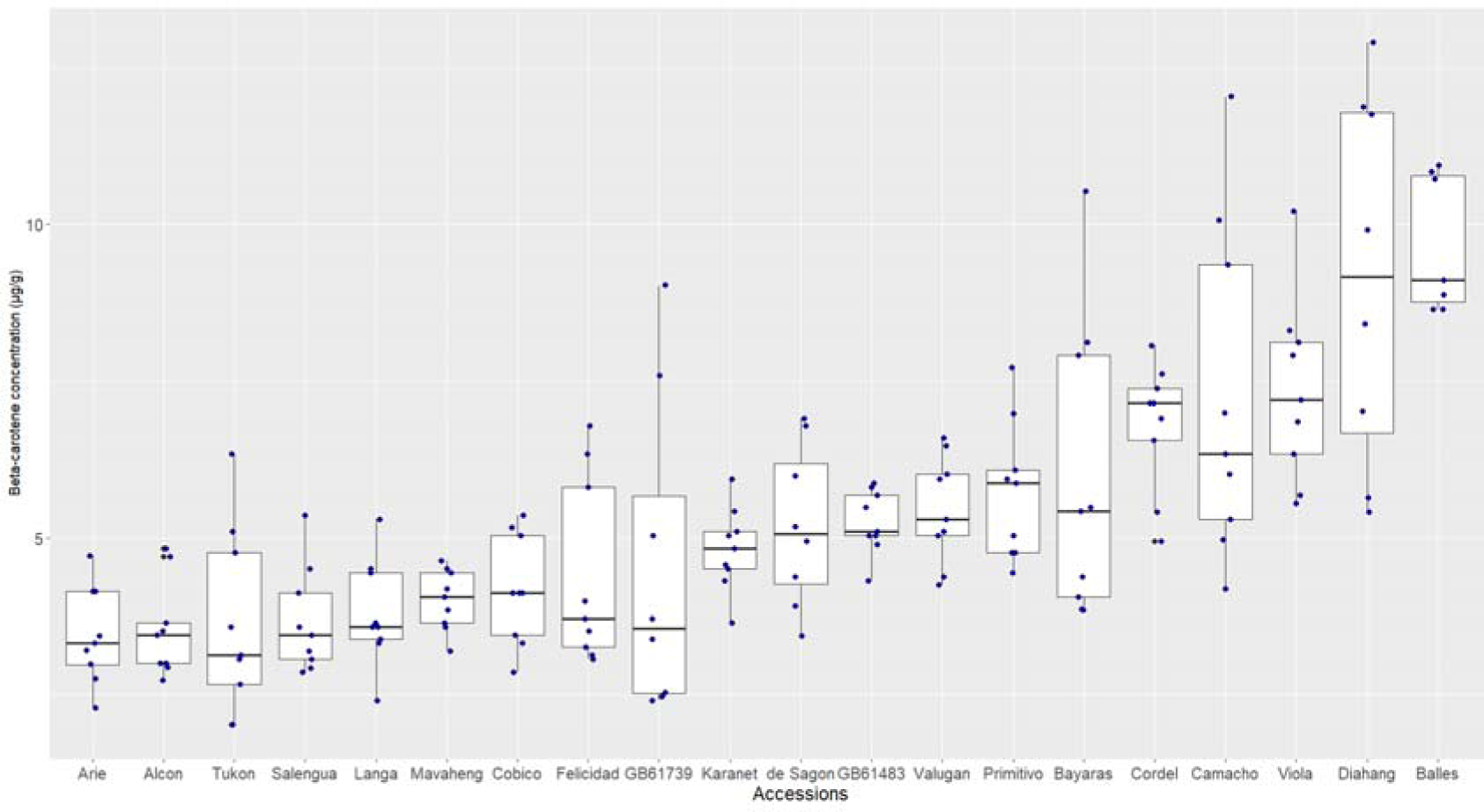
β-carotene concentrations of 20 foxtail millet accessions (in µg/g).

### Correlations among the inorganic nutrients

Correlation analysis of the micronutrients, including N and β-carotene, using Pearson’s rank correlation, identified a number of significant correlations (Fig. 7). There was a significant, positive correlation between Ca and Zn (*R* = 0.66, *P* < 0.001), between Zn and Cu (*R* = 0.43; *P* < 0.001), between Mg and Cu (*R* = 0.40; *P* <0.001), and a negative correlation between Mg and N (*R* = –0.43, *P* < 0.001). Other correlations are shown in Fig. 7.

**Fig. 7.**
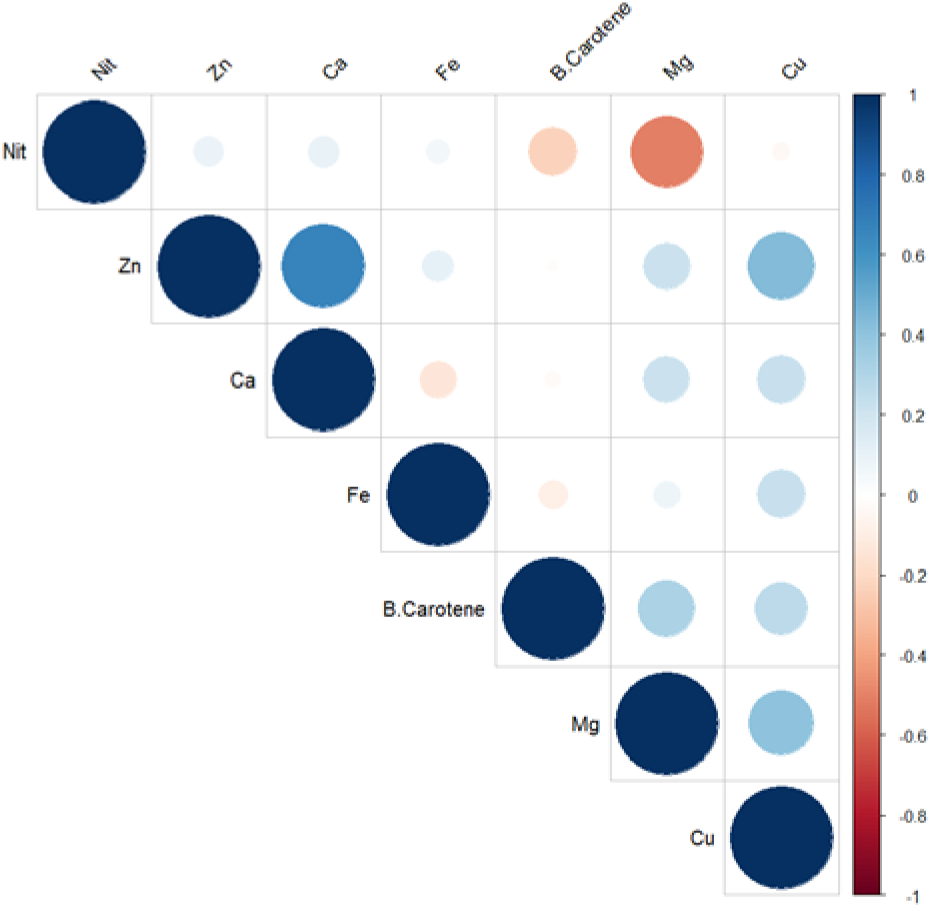
Pearson correlations among the micronutrients.

### Phenolic compound content

Flour samples of the foxtail millet accessions were screened for 20 phenolic compounds, with significant differences (*P* < 0.001) among the 20 accessions being found for 12 phenolic compounds (Suppl. Table 11).

Catechin, kaempferol, myricetin, and quercetin are phenolic compounds classified as flavonoids. Quercetin was present at a low mean concentration of 0.10 µg/g, with most of the quercetin being present in its glycosylated state, as quercetin glycoside (mean 0.31 µg/g). Myricetin was synthesized at an average concentration of 0.32 µg/g. Kaempferol and its glycoside product, kaempferol glycoside, were both present, with mean concentrations of 1.51 and 0.84 µg/g, respectively (Fig. 8). The flavonoid catechin was found to be the highest phenolic synthesized, with an average value of 4.50 µg/g (Fig. 8; Suppl. Table 11). Syringic acid and ferulic acid were present at high levels, with average values of 4.39 and 2.97 µg/g, respectively (Fig. 8; Suppl. Table 11).

**Fig. 8.**
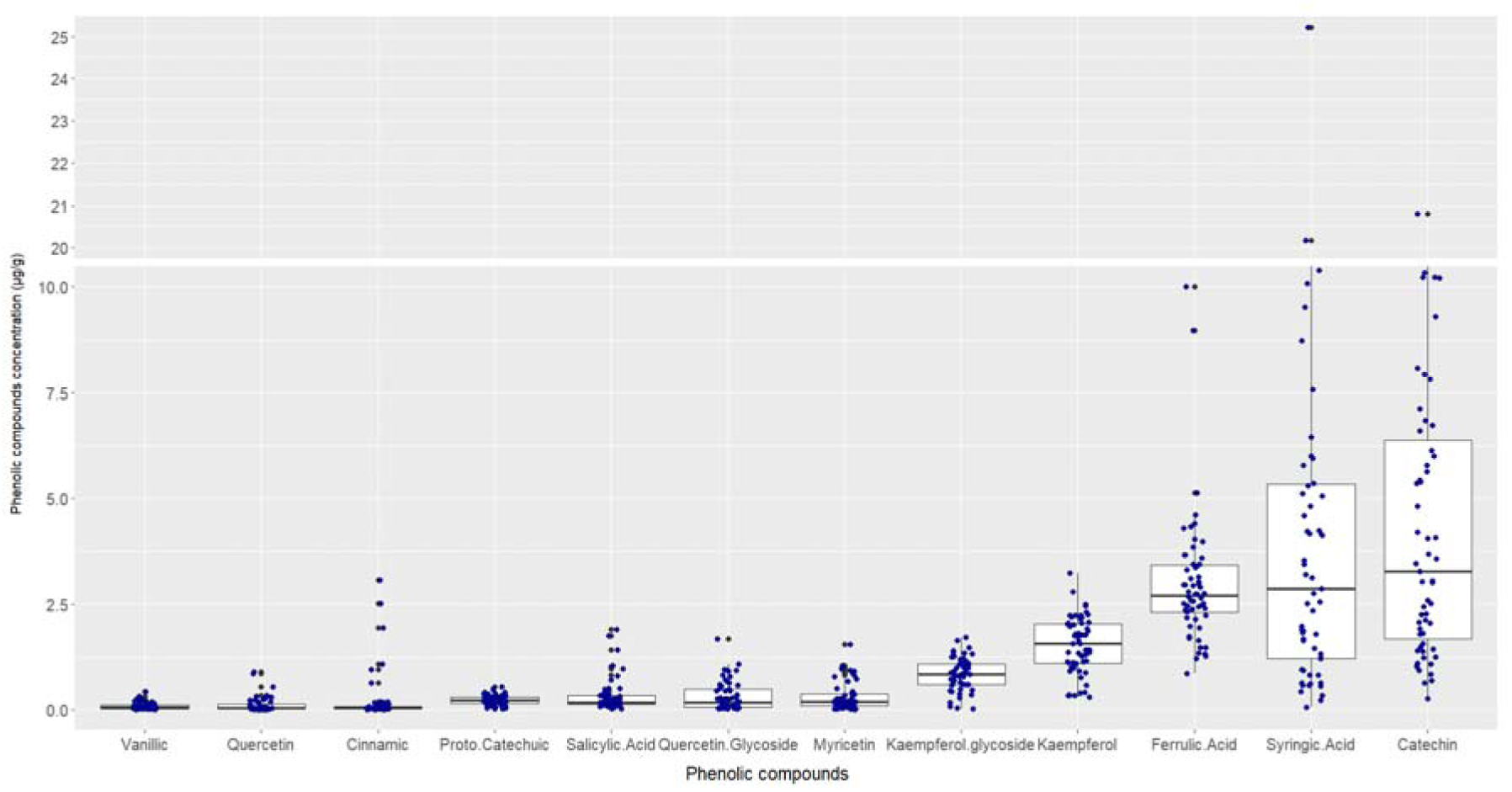
Variations in the content of 12 phenolic compounds detected across all foxtail millet accessions. (Note that the plot has been truncated).

GB61483 was found to produce the highest levels of catechin, syringic and ferulic acid, with average concentrations of 20.61, 17.12, and 2.84 µg/g, respectively (Suppl. Table 11).

A Pearson’s correlation analysis was undertaken between the phenolic compounds (Fig. 9B). The highest positive correlation was seen between catechin and ferulic acid (*R* = 0.90, *P*<0.001), followed by cinnamic acid and quercetin (*R* = 0.86, *P*<0.001). Strong correlations were also found between ferulic acid and kaempferol (*R* = 0.65; *P* < 0.001), ferulic acid and syringic acid (*R* = 0.62; *P* < 0.001) and ferulic acid and quercetin glycoside (*R* = 0.59; *P* < 0.001), with strong positive correlations also seen between catechin and kaempferol (*R* = 0.68; *P* < 0.001) catechin and syringic acid (*R* = 0.54; *P* < 0.001) and catechin and quercetin glycoside (*R* = 0.70; *P*<0.001). The biochemical relationship between these phenolic compounds is shown in Fig. 9A.

**Fig. 9.**
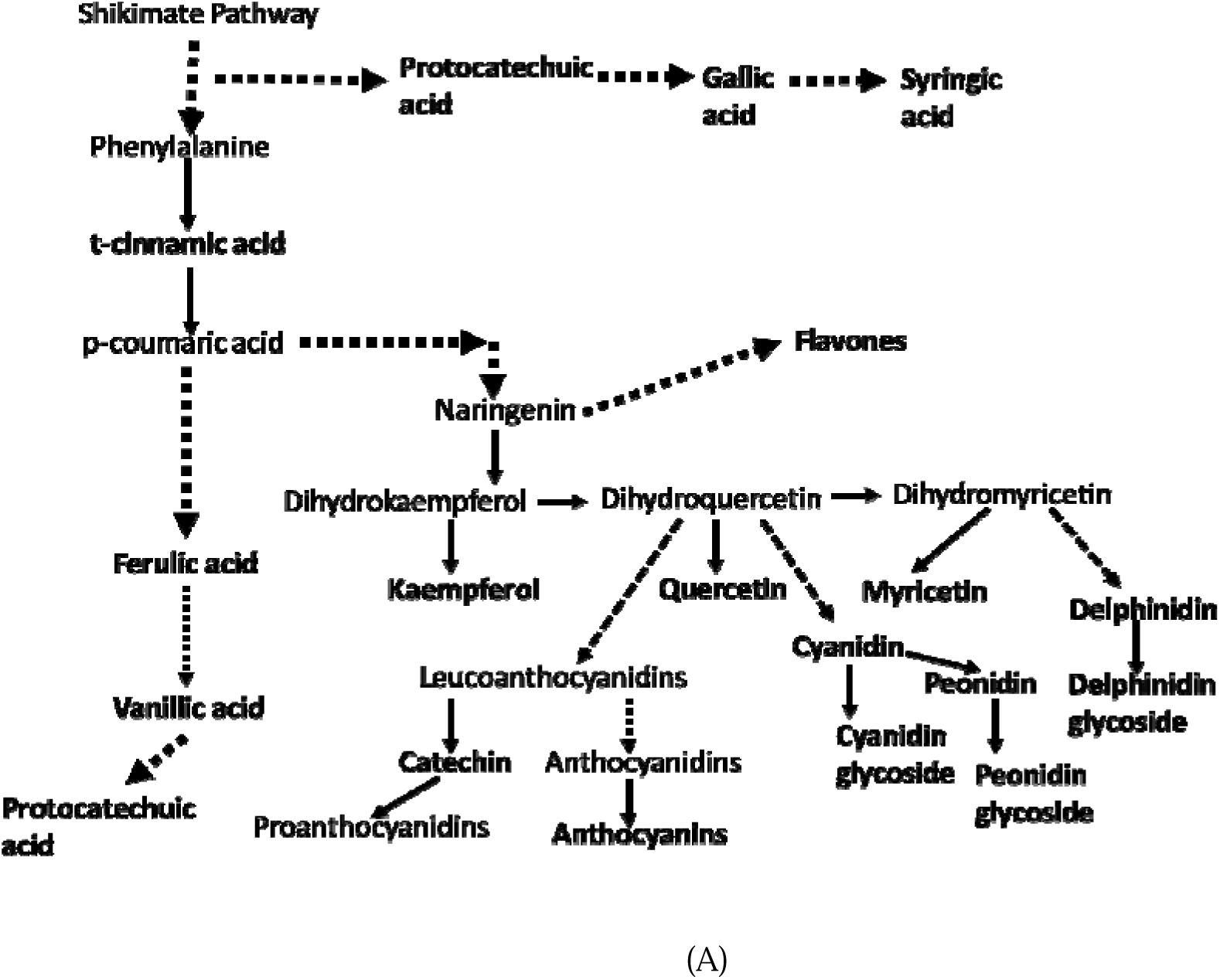
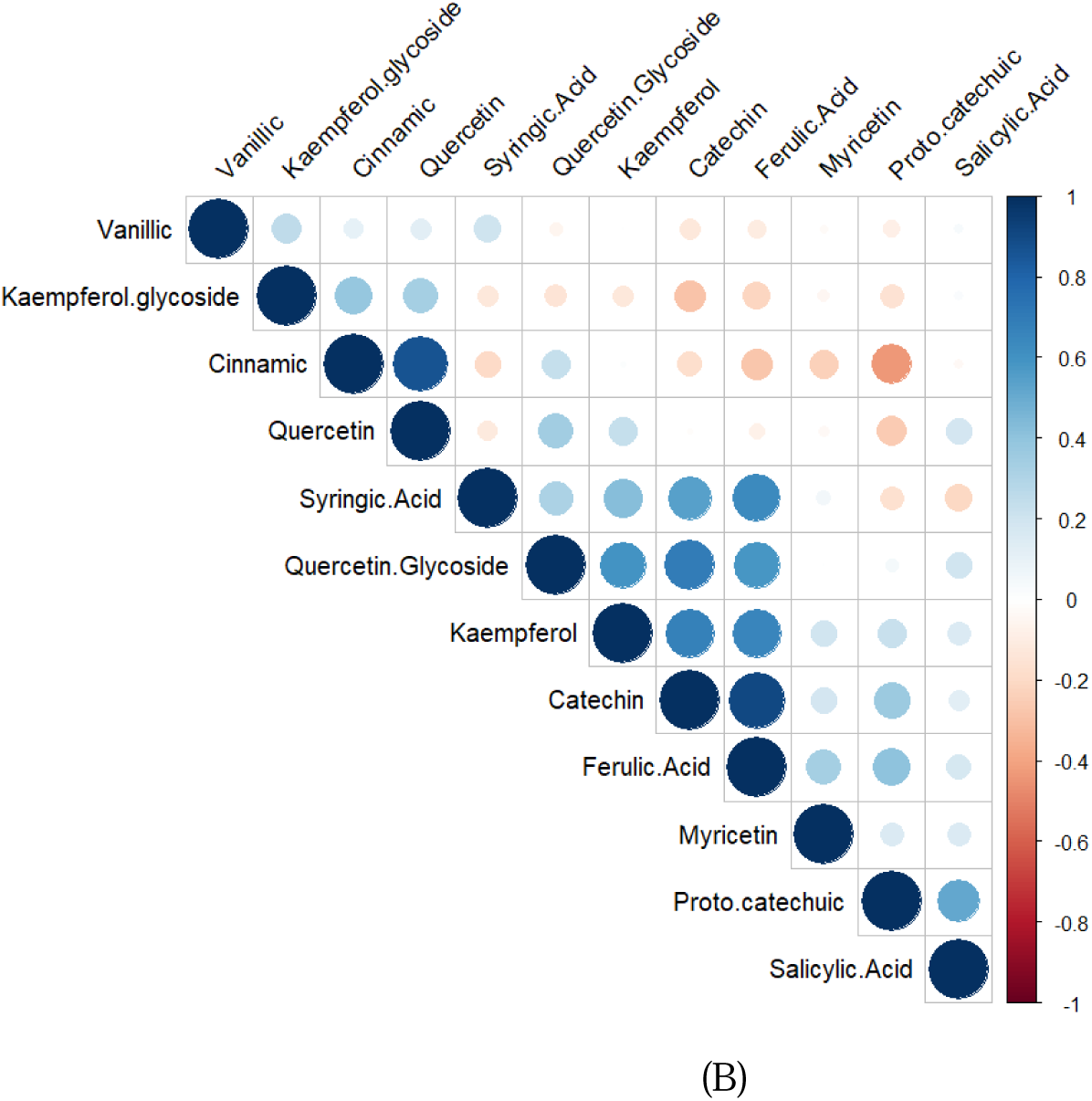
(A) Biochemical pathway relationship between phenolic compounds assessed in this study [adapted from 45;46]. Dashed lines indicate multi-step reactions. (B) Correlations between the levels of phenolic compounds detected in this study.

### Whole genome sequencing analysis

We wanted to determine the genetic diversity within a core set of local foxtail millet accessions. Eight accessions (Table 5) were, therefore, selected for whole genome sequencing (WGS), using Illumina NovaSeq paired end (PE150) technology, at a 25× coverage. Bayaras and GB61438 were selected for sequencing as they were found to accumulate high levels of Zn and Fe, respectively. For comparison, we sequenced the lowest Fe- and Zn-accumulating lines, Felicidad and GB61739, respectively. Palacu and Ukig were included as they are geographically different from the rest of the lines. Mavaheng and de Sagon were randomly selected to represent reddish brown and very light brown accessions, respectively, from Batanes. Note that both GB* accessions were of unknown origins, therefore, their genetic relatedness to the other accessions would be of interest.

A total of 862,100,516 raw reads were generated across all eight accessions (Suppl. Table 12). Data quality metrics Q20 and Q30 showed percentages of more than 97% and 93%, respectively, suggesting relatively high-quality datasets. Quality assessment of the reads using FASTQC indicated no residual adapter sequences, over-represented sequences, or low-quality bases. Thus, no further pre-processing step was performed.

Mapping of the reads showed more than 99% alignment to the Yugu1 reference genome. Variant calling using GATK4 (HaplotypeCaller; by-sample, in VCF mode) identified more than 3 Mb of unfiltered variants for each accession. Implementation of filtering parameters revealed that Ukig and GB61739 have the highest and lowest number of variants, with 968,517 and 826,818, respectively (Table 4). There were also considerable numbers of Indels, with Ukig having the highest number with 82,225 Indels and GB61438 the lowest, with 70,052 Indels. Variant rate calling indicated that Ukig contained 2.39 variants for every 1 kb (or 1 in every 418 bases), while GB61438 had 2.04 variants per kb (or 1 in every 489 bases).

### Population structure and phylogeny

We identified 2,046,040 SNPs in GVCF mode (for multi-sample cohorts) across all eight accessions after implementing several filtering parameters with VCFtools (Indels and monomorphic SNPs removed; see Materials and Method). Using the genome-wide SNP data as an input, the population structure was inferred viz. fastStructure across varying number of populations (K). The distruct plot consistently partitioned the accessions into two unadmixed groups at varying K values (K = 2 to 5) (Fig. 10A). The first group includes the very light brown accessions (Bayaras, Felicidad, de Sagon, and GB61739); the other, the light brown (Palacu and Ukig) and reddish-brown accessions (GB61438 and Mavaheng).

**Fig. 10.**
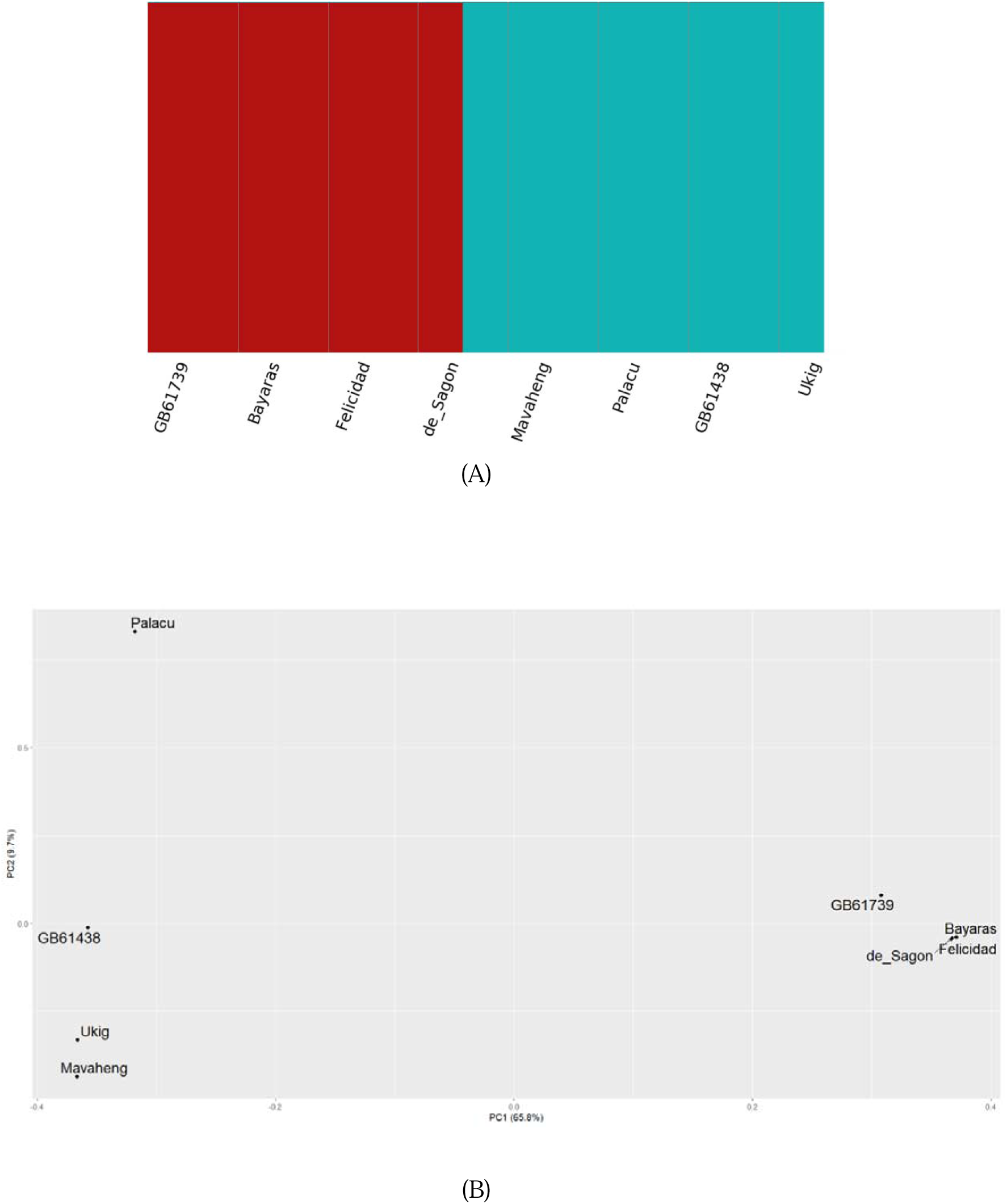
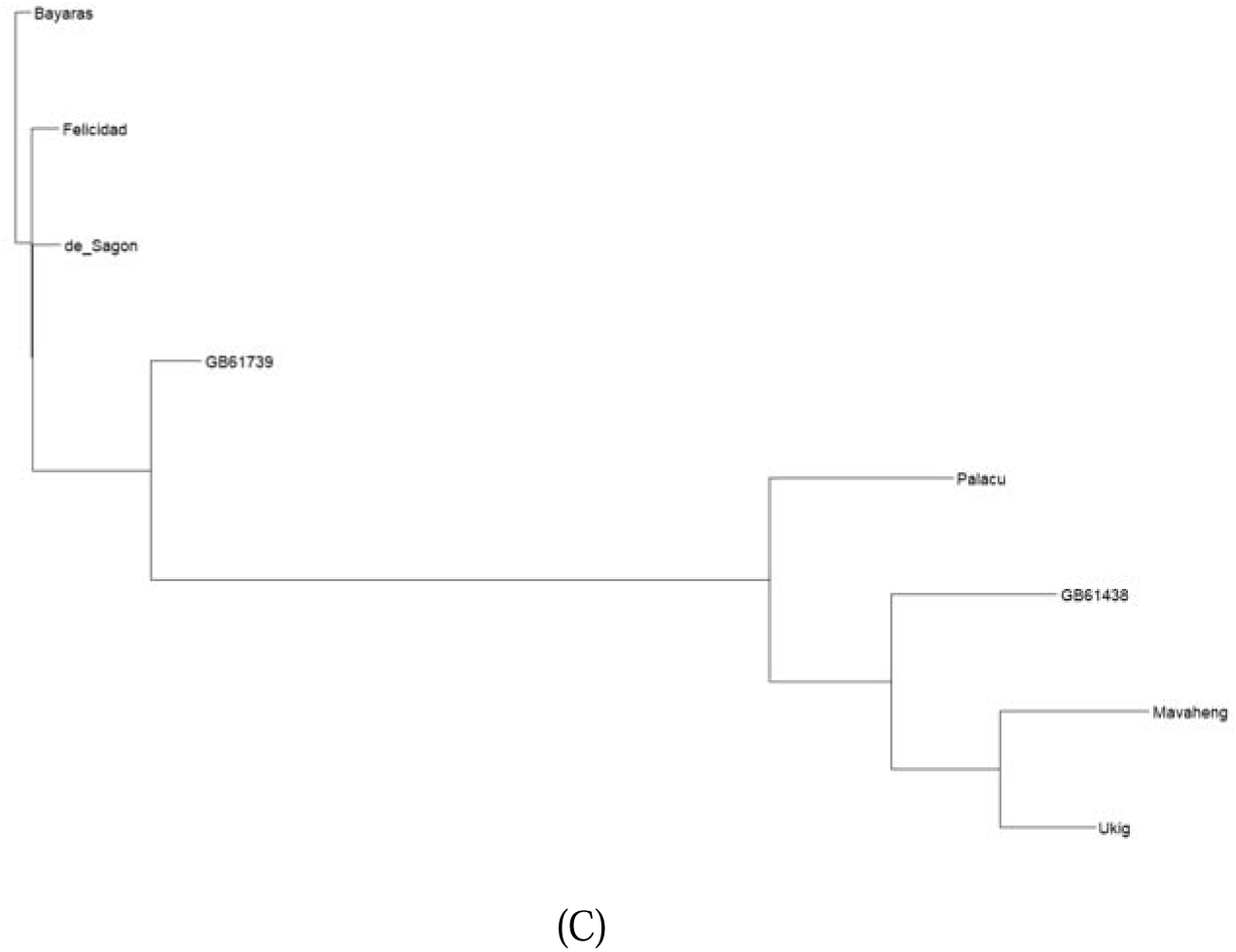
(A) Structure analysis assuming K = 2; (B) PCA; (C) Phylogenetic tree implementing Neighbor–Joining algorithm in VCF kit.

The finding that the accessions were partitioned into two groups was consistent with the PCA (Fig. 10C) in which PC1 (x-axis) explained the majority of the variation (65.8%) among the eight varieties, segregating them into two groups (Fig. 10C). PC2 (y-axis) explained 9.7% of the variation.

To further determine the evolutionary relationship among the sequenced accessions we generated a phylogenetic tree (Fig. 10D). Relative to their earliest common ancestor, the accessions are monophyletic, with the very light brown accessions Bayaras, Felicidad, de Sagon, and GB61739 emerging earlier than the light brown (Palacu and Ukig) and reddish-brown accessions (GB61438 and Mavaheng). As shown by the short branch lengths (Fig. 10C), Felicidad, de Sagon and Bayaras are apparently closely related, and showed limited divergence. This is further confirmed by the PCA, in which all three accessions are clustered contiguously. As the GB* accessions are of unknown origin, we can infer from these analyses of genetic relatedness that GB61438 is genotypically close to Ukig, a Bicol accession, while GB61739 sits close to the Batanes accessions Felicidad and Bayaras.

### Zinc Iron Permease (ZIP) transporter

We were particularly interested in elucidating the molecular mechanism behind Zn and Fe uptake, as these are nutritionally limited in the Philippines. The lack of both minerals being a chronic problem in the country. Both metals are actively accumulated by transporters from the ZIP transport family of proteins. The EBI Interpro database showed that Pfam code PF02535 is associated with ZIP transporters (integrated in Interpro, IPR003689). This family was mainly contributed from *Arabidopsis thaliana* [47]. To identify putative ZIP sequences, we searched Ensembl Plant (*Setaria italica* ref Yugu1) with the code PF02535. Results returned 17 hits of genes containing this protein domain. This is comparable to *A. thaliana* and rice which harbour 15 and 17 ZIP transporters, respectively [48]. However, recent interrogation of Ensembl Plants revealed 20 genes in indica and 15 in japonica rice. The 17 ZIP genes were scattered across the foxtail millet reference genome of var. Yugu1, being located on chromosomes I (2 genes), II, III (3 genes per chromosome), IV, VI (3 genes per chromosome), VII (3 genes), and IX (4 genes).

Phylogenetic analysis of the translated amino acid sequences of the ZIP transporters from foxtail, *A. thaliana* and rice (japonica) indicated a closer relationship between foxtail and rice ZIP proteins (Fig. 11). This is expected as both are cereal monocots, while *A. thaliana* is a dicot. In most cases, for every single foxtail ZIP protein there was a corresponding rice ZIP protein, suggesting parallel descent from a common ancestral gene. In a few cases, rice ZIP proteins were found associated with two foxtail millet ZIP proteins. Example, rice protein ZIP9 was found linked with the two foxtail millet ZIP proteins, KQL15068 (Gene: SETIT_024505mg) and KQK98376 (Gene: SETIT_010745mg). These two are said to be paralogs (indicated by blue arrows in Fig. 11). On the other hand, the paralogs KQL29912 (Gene: SETIT_016756mg) and KQK86158 (Gene: SETIT_034876mg) were found associated with a single rice gene, OsZIP (Os05t0316100-01).

**Fig. 11.**
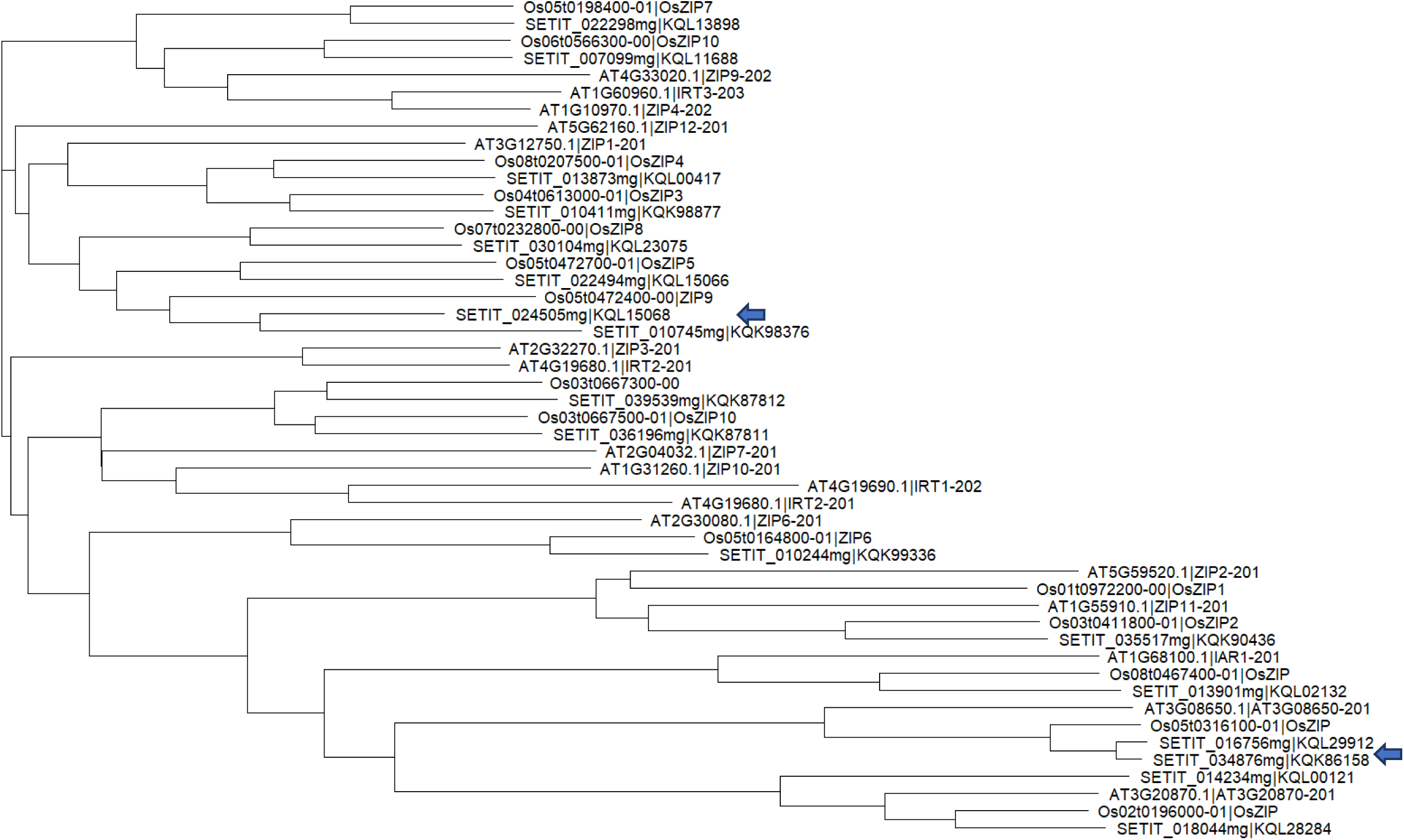
Phylogenetic tree of ZIP transporter of foxtail millet (*Setaria italica*; prefixed with SETIT), rice (*Oryza sativa*; prefixed with Os), and *Arabidopsis thaliana*; prefixed with AT). Figure generated using VCF-kit in R. Blue arrows indicate location of foxtail ZIP protein paralogs.

As we found that the mineral levels in foxtail millet are comparable, or even higher than those found in conventional rice varieties, we further scrutinized the foxtail millet ZIP transporter gene sequences. Using SNPEff, we detected several SNPs found within 14 of the 17 ZIP transporter genes. Most of these SNPs were located in the 5’ upstream region (109 SNPs), in intron (174 SNPs), and in 3’ downstream regions (284 SNPs) (Suppl. Table 13) suggesting that the SNPs may mainly affect gene regulation rather than altering the function of the protein through changes in the DNA sequence of the coding region.

## Discussion

The Philippines has chronic food insecurity and malnutrition problems, in particular Zn and Fe deficiencies. Its government has proactively initiated the identification of alternative sources of nutrients, including biofortification of food products, particularly rice, to address these problems. Therefore, in this study, we have investigated the nutritional potential of foxtail millet, an heirloom crop, as an alternative source of nutrients. We collected several accessions of this crop, grown at dispersed locations within the country. The levels of selected micronutrients, N (as a proxy to crude protein), β-carotene and phenolic compounds were quantified. The levels of protein, measured as a proxy of N levels, found in this collection of foxtail millet accessions (5.63 to 13.38 g/100g) were comparable to levels found in other accessions of foxtail millet collected from Asian countries, including Bangladesh and India, where protein levels ranged from approx. 11.65 g/100g [7] to 12.3 g/100g [9; 10], respectively. These protein levels are higher than those reported in brown rice by the USDA (∼2.7 g/100g; [49]).

While Zn concentrations in the soil lie on the medium scale, the foxtail millet accessions were able to accumulate high levels of this metal in the grain. This would suggest a greater ability of foxtail millet to transport Zn from the soil and store it at higher levels in the grain. Furthermore, most of the foxtail millet accessions demonstrated Zn levels higher than the recommended 40 mg/kg of Zn in flour [50], and higher than those reported in rice. Brown rice was reported to contain 2 mg/100g (or 20 mg/ kg) [51] of Zn; while golden rice (GR) and the near isogenic control PSBRc82 contained Zn levels of 2.31 mg/100g (or 23.1 mg/kg dry basis (DB)) and 2.19 mg/100g (or 21.9 mg/kg DB), respectively [52]. The levels in foxtail millet are also far higher than the Zn levels found in tef (*Eragrostis tef*), another neglected and underutilized crop, where Zn levels range from 14.8 to 29.2 mg/kg [28]. However, it must be remembered that direct comparisons with the literature have to be considered with caution, as these crops were not grown in the same season and experimentation site.

Mean Fe concentrations in foxtail millet were comparable with or even higher than those reported in rice, which contains on average 40 mg/kg DB (for GR) and 46 mg/kg (for PSBRc82) [52]. The high concentration of Fe in the foxtail millet grains may be ascribed to the high soil concentration of Fe at the field site (Table 3). Mg, Ca and Cu levels were found to be comparable to previous reports [7]. These metals were found to have concentrations of around 454, 470 and 5.8 mg/kg, respectively.

Foxtail millet therefore presents as a very promising alternative source of micronutrients, in particular Zn, and an ideal crop to supplement the Filipino diet and address Zn deficiency. Adults require 8–11 mg Zn/ day while lactating and pregnant women require 11–13 mg [53]. This means, ∼200 grams of foxtail millet can satisfy the daily Recommended Nutrient Intake. It is interesting to note that while there was a minimal input applied (i.e., no fertilizer added, once-a-week watering), the crop exhibited high levels of nutrients, particularly protein, Zn and Fe. The Nitrogen use efficiency (NUE) of this crop would be an interesting subject of further inquiry. Furthermore, this crop may be a potential source of genetic features for the enhancement of the nutritional value of other staple cereal crops, viz. genetic modification.

Positive correlations were seen between the levels of Zn with both Ca and Cu, and between Mg and Cu, while a negative correlation was found between Mg and N. A study in spring wheat found that slightly acidic soils increased the level of ammonium and hydrogen ions, which may in turn have a competitive effect on the uptake of Mg [54]. The slightly acidic profile of the field sites (pH 5.97 to 6.12; Table 3) may explain the negative correlation seen in this study between N and Mg.

With regards to Zn and Ca, an antagonistic interaction has been shown between these metals which leads to an ameliorative effect of Ca on Zn deficiency [55]. However, in foxtail millet a strong positive correlation was found between Zn and Ca. Assimilation of both metals synergistically suggests that there could be other factors in foxtail millet that are influencing uptake of these metals.

β-carotene has been a major target for nutritional improvement, especially in developing countries. In rice, conventional varieties show levels of β-carotene below the limit of quantification (LOQ) [52]. In the genetically engineered GR, total carotenoid levels in the of range 3.5−10.9 mg/kg DB (mean 5.88 mg/kg DB) were measured, while-trans-β-carotene levels ranged from 1.96−7.31 mg/kg (mean 3.57 mg/kg) [52], which is lower than the β-carotene concentrations found in most of the foxtail accessions. However, this study used 3-month-old endosperms. An earlier study, using freshly harvested rice seed, found up to 36.7 µg/g of carotenoid, 84.1% of which was β-carotene, in GR [56]. Carotenoids are known to degrade with storage, and their stability is influenced by many other external factors [57]. The finding that foxtail millet is loaded with β-carotene provides an opportunity for future research, and the possibility of supplementing rice grain with foxtail millet seed as a pro-Vitamin A source in the diet. A study of the carotenoids lutein and zeaxanthin (β-carotene was not measured) reported 13.58 mg/kg of lutein and 3.77 mg/kg of zeaxanthin in the foxtail millet variety Jingu21, and 0.81 mg/kg of lutein and 0.22 mg/kg of zeaxathin in the variety Zhisheng [58].

Twelve phenolic compounds were detected in the foxtail millet accessions (Fig. 8). However, delphinidin, cyanidin and peonidin, and their glycosides, the anthocyanins which confer colour in plants and cereal grain, were not detected. Eight of the compounds were detected at relatively low levels (Fig. 8). However, kaempferol, ferulic acid, syringic acid and catechin were present at higher levels and exhibited considerable variation between the 20 accessions. Strong positive correlations were seen between these four compounds. The relationship between catechin and kaempferol is apparent, both being derived from naringenin, however ferulic acid and syringic acid sit on different arms of the shikimate pathway (Fig. 9A).

In rice, high catechin levels has been found to be associated with a low glycaemic index and antioxidant properties [59]. Studies on green tea (*Camellia sinensis*) showed that catechin has a hypocholesterolaemic effect and anti-inflammatory properties [60]. Syringic acid was previously shown to confer therapeutic effects on diabetes, cancer and cardiovascular diseases, having antimicrobial, antioxidant and anti-inflammatory properties (reviewed in [61]). Food rich in kaempferol have also been associated with a decreased risk of developing cancers such as skin and colon cancers [62].

Analysis of the genetic variation in eight of the foxtail millet accessions divided the accessions into two groups, suggesting a rather modest diversity among the local accessions. Bayaras, Felicidad, de Sagon and GB61739 clearly clustered together, having one inferred ancestral population unadmixed. Mavaheng, Palacu, Ukig and GB61438 also appear to have one common unadmixed ancestry. Phylogenetic analysis showed that the accessions are monophyletic. This finding is consistent with a recent paper in which foxtail millet accessions are monophyletic with respect to the wild ancestor, green foxtail [22]. Variant calling analysis showed that Ukig carries more variants compared to the other accessions. Ukig is cultivated on an island of the Bicol Region called Catanduanes (see Fig. 1). Phenotypically, Palacu was observed to exhibit a shorter life cycle, flowering ∼50 DAS as compared to other accessions which flowered from 56 to 92 DAS. Furthermore, Palacu is cultivated in Cagayan, a different eco-geographical location relative to the other accessions (Table 2).

As Zn and Fe are actively accumulated in plants by transporters from the ZIP transport family of proteins, we screened the *S. italica* reference genome Yugu1 with the Pfam code PF02535 (integrated in Interpro IPR003689). This search identified 17 ZIP transporter protein domains in the *S. italica* reference genome Yugu1. This number of putative ZIP transporters in comparable to rice, a diploid cereal with 17 putative ZIP transporters. The cereal tef has been reported to have 32 putative ZIP transporters [28] and bread wheat 60 ZIP proteins [34], however these cereals are tetraploids and hexaploids, respectively. Several SNP variants were found among the foxtail millet accessions, located primarily within 3’ and 5’ regulatory regions and introns. This would suggest that variation in Zn and Fe levels maybe due to variation in the regulation of ZIP transporter levels, rather than in protein structural differences. Many studies have reported the role of cis and trans elements in gene regulation [63; 64], and promoter sequences and cis-regulatory elements have been shown to influence iron response mechanisms in rice [65]. However, transcription can also be influenced by the environment [66; 67]. Therefore, the variation in Zn and Fe accumulation observed between foxtail millet accessions may be largely ascribed to the interaction of the environment (e.g., availability of the metals in the soil, pH) and the genotype (e.g., variations in regulatory elements) (G × E), and is needing of further study. Future breeding of foxtail millet for enhanced nutritional value would benefit from the identification of the underlying genes, gene-specific SNPs and haplotypes associated with Zn and Fe accumulation, as well as the other beneficial nutritional compounds identified in this study. While such studies would require analysis of a larger population of foxtail millet accessions the work reported here demonstrates the considerable phenotypic and genetic variation that exists in foxtail millet for nutritionally valuable traits, supporting further analyses.

## Conclusion

In this study we report the potentials of foxtail millet, a neglected and under-utilised cereal crop, for addressing the issues associated with the double-burden of malnutrition. We have demonstrated the potential of foxtail millet to accumulate high levels of Zn, even in soils that have only an average level of this mineral, along with high levels of phenolic compounds such as catechin, which in rice has been shown to reduce glycaemic index, lowering the risk of type-2 diabetes. The local foxtail millet accessions found across the Philippines represent a potential alternative source of nutrients for humans (and animals). Due to its modest cultivation and diversity, we wish to highlight the need to ensure the conservation of this cereal as it may contribute in addressing malnutrition particularly Zn, Fe, and β-carotene.

## Data availability

All sequencing datasets are available at the ENA under ArrayExpress accession E-MTAB-12658 and E-MTAB-13027.

## Ethical standards on the use of materials

We adhered to the highest ethical standards on the use of the biological materials. No ethical implications are associated on the use of these plant materials as foxtail millet is not a protected species in the Philippines.

## Authors’ Contributions

NCE conceptualized the project, sought project funding, performed statistical and bioinformatics analyses, and wrote the initial draft of the manuscript. JAR undertook data collection and management of the field trials. DEC performed chemical analysis of the flour. RA undertook molecular biology work and bioinformatics work. NMO performed analysis of the soil samples. HJ undertook phenolics and β-carotene analyses. LB managed the analysis of the samples at NIAB, wrote the manuscript. ED managed the lab and field activities. All authors reviewed and approved of the publication of this work.

## Acknowledgement

We thank the Ivatans of Batanes, classified as Indigenous People of the Philippines, for allowing the use of their materials for this research. The authors thank Dr. Raquel Salingay, Rafael Magallanes, Leila Baculi and Melvin Allan M. Hipol for donating millet seeds used in this study. We also thank North Central Regional Plant Introduction Station (NCRPIS) for providing the Taiwanese germplasm. We thank the staff of IPB Labs (Juliet Welgas, Rolyn Gonzales, Edgardo Reyes, Elenita R. Castillo, Nistan Jay V. Hadjinulla, and Lizk Gil P. Villegas) and ICropS (Dr. Tonette Laude) for their assistance in the greenhouse activities, field trials, and chemical analysis. The lead author, NCE, would like to thank Cambridge Global Challenges and Global Food Security of the University of Cambridge, UK for awarding travel funds to cover transportation to and from the Philippines to initiate this project. The phenolic analyses were funded by the UK Biotechnology and Biological Sciences Research Council (BBSRC) project BB/T008873/1. This work was funded by the UP System Enhanced Creative Work and Research Grant (ECWRG-2021-1-10R).

## Abbreviations

ZIP: Zinc/Iron Permease
BLAST: Basic Local Alignment Search Tool
SA: Syringic Acid
DS: Dry Season
WS: Wet Season
MSA: Multiple Sequence Alignment

## Supplementary Tables

Suppl. Table 1. Percentage (or g/100g) means and standard deviations of ash content among the accessions

Suppl. Table 2. Total N and Crude Protein contents (raw data) across all accessions

Suppl. Table 3. Zn contents of the different accessions (raw data)

Suppl. Table 4. Analysis of Zn content using Tukey’s Honesty Significant Difference

Suppl. Table 5. Fe contents of the different accessions (raw data)

Suppl. Table 6. Mg contents of the different accessions (raw data)

Suppl. Table 7. Ca contents of the different accessions (raw data)

Suppl. Table 8. Cu contents of the different accessions (raw data)

Suppl. Table 9. Beta carotene contents of the different accessions (raw data)

Suppl. Table 10. Analysis of β-carotene content using Tukey’s HSD

Suppl. Table 11. Phenolics contents of the different accessions (raw data)

Suppl. Table 12. Data quality information of the raw reads

Suppl. Table 13. Information of SNPs co-localizing with foxtail millet ZIP genes

## References

1. Diao X, Jia G. Origin and Domestication of Foxtail Millet. In: Genetics and Genomics of Setaria. Cham: Springer International Publishing; 2017. p. 61–72.

2. Zhang G, Liu X, Quan Z, Cheng S, Xu X, Pan S, et al. Genome sequence of foxtail millet (*Setaria italica*) provides insights into grass evolution and biofuel potential. Nat Biotechnol. 2012;30(6):549–54. Available from: http://dx.doi.org/10.1038/nbt.2195

3. Doust AN, Km Devos JL. Foxtail millet: a sequence-driven grass model system. Plant Physiol. 2009; 149:137–41. https://doi.org/10.1104/pp.108.129627

4. de Wet JMJ, Oestry-Stidd LL, Cubero JI. Origins and evolution of foxtail millets (*Setaria italica*). J Agric Tradit Bot Appl. 1979;26(1):53–64. Available from: http://dx.doi.org/10.3406/jatba.1979.3783

5. Barton L, Newsome SD, Chen F-H, Wang H, Guilderson TP, Bettinger RL. Agricultural origins and the isotopic identity of domestication in northern China. Proc Natl Acad Sci U S A. 2009;106(14):5523–8. Available from: http://dx.doi.org/10.1073/pnas.080996

6. Verma S, Srivastava S, Tiwari N. Comparative study on nutritional and sensory quality of barnyard and foxtail millet food products with traditional rice products. J Food Sci Technol. 201552(8):5147–55. doi: 10.1007/s13197-014-1617-y.

7. Abedin MJ, Abdullah ATM, Satter MA, Farzana T. Physical, functional, nutritional and antioxidant properties of foxtail millet in Bangladesh. Heliyon. 2022; 8(10):e11186. Available from: http://dx.doi.org/10.1016/j.heliyon.2022.e11186.

8. FAO. 2010. Available from http://www.fao.org/ag/agn/nutrition/phl_en.stm.

9. . Kumari R, Dikshit N, Sharma D, Bhat KV. Analysis of molecular genetic diversity in a representative collection of foxtail millet [*Setaria italica* (L.) P. Beauv.] from different agro-ecological regions of India. Physiol Mol Biol Plants. 2011;17(4):363–7.

10. Kamatar MY, Brunda SM, Sanjeevsingh R, Sowmya HH, Giridhar G, Ramaling H, et al. Nutritional composition of seventy five elite germplasm of Foxtail millet (*Setaria italica*). Int J Eng Res Technol (Ahmedabad). 2015;V4(04). Available from: http://dx.doi.org/10.17577/ijertv4is040075

11. Fukunaga K, Kato M. Geographical variation of nuclear genome RFLPs and genetic differentiation in foxtail millet, Setaria italica (L.) P. Beauv. Genet Resour Crop Evol. 2002; 49:95–101.

12. Wei L, Hui Z, Yong-fang W, Hai-quan L, Xian-mm D. Assessment of Genetic Relationship of Foxtail Millet with Its Wild Ancestor and Close Relatives by ISSR Markers. J Integr Agric 2012; 11(4): 556–566. Available from https://doi.org/10.1016/S2095-3119(12)60042-2

13. Yadav RK, Adhikari AR, Gautam S, Ghimire KH, Dhakal R. Diversity sourcing of foxtail millet through diversity assessment and on-farm evaluation. Cogent Food Agric. 2018;4(1):1482607. Available from: http://dx.doi.org/10.1080/23311932.2018.1482607

14. Zhang G, Liu X, Quan Z, Cheng S, Xu X, Pan S, et al. Genome sequence of foxtail millet (*Setaria italica*) provides insights into grass evolution and biofuel potential. Nat Biotechnol. 2012;30(6):549–54. Available from: http://dx.doi.org/10.1038/nbt.2195

15. Li C, Wang G, Li H, Wang G, Ma J, Zhao X, et al. High-depth resequencing of 312 accessions reveals the local adaptation of foxtail millet. Züchter Genet Breed Res. 2021;134(5):1303–17. Available from: http://dx.doi.org/10.1007/s00122-020-03760-4

16. Salingay, Raquel O., Agripina R. Aradilla, Nenita B. Baldo, Ma. Stella M. Paulican. 2021. Ethno-production and Utilization Practices of Foxtail Millet (*Setaria italica* (L.) P. Beauv) in Northern Mindanao, Philippines. International Journal of Academic and Applied Research (IJAAR). 5(11)45–53.

17. Sakamoto S. Origin and Dispersal of Common Millet and Foxtail Millet. JARQ. 1987;21(2).

18. Nasu H, Momohara A, Yasuda Y, He J. The occurrence and identification of *Setaria italica* (L.) P. Beauv. (foxtail millet) grains from the Chengtoushan site (ca. 5800 cal B.P.) in central China, with reference to the domestication centre in Asia. Veg Hist Archaeobot. 2007;16(6):481–94. Available from: http://dx.doi.org/10.1007/s00334-006-0068-4

19. . Ceasar SA, Ramakrishnan M, Vinod KK, Roch GV, Upadhyaya HD, Baker A, et al. Phenotypic responses of foxtail millet (*Setaria italica*) genotypes to phosphate supply under greenhouse and natural field conditions. PLoS One. 2020;15(6):e0233896. Available from: http://dx.doi.org/10.1371/journal.pone.0233896

20. Edmondson, RN. Multi-level Block Designs for Comparative Experiments. JABES. 2020; 25, 500–522. doi:10.1007/s13253-020-00416-0

21. Nadeem F, Ahmad Z, Ul HM, Wang R, Diao X, Li X. Adaptation of Foxtail Millet (*Setaria italica* L.) to Abiotic Stresses: A Special Perspective of Responses to Nitrogen and Phosphate Limitations. Front. Plant Sci. 2020; 11:187. doi: 10.3389/fpls.2020.00187

22. Hunt HV, Przelomska NAS, Campana MG, Cockram J, Bligh HFJ, Kneale CJ, et al. Population genomic structure of Eurasian and African foxtail millet landrace accessions inferred from genotyping-by-sequencing. Plant Genome. 2021;14(1):e20081. Available from: http://dx.doi.org/10.1002/tpg2.20081

23. Peech M, Alexander LT, Dean LA, Reed JF. Methods of soil analysis for soil fertility investigation. USDA Circ. 1947.

24. Walkley A, Black CA. An examination of Degtiareff method for determination of soil organic matter and a proposal modification of the chromic acid titration method. Soil Sci. 1934;37:29–35.

25. Bray R, Kurtz LT. Determination of total, organic, and available forms of phosphorus in soils. Soil Sci. 1945;59:39–45.

26. Lindsay WL, Norvell WA. Development of a DTPA soil test for zinc, iron, manganese, and copper. Soil Sci Soc Am J. 1978;42(3):421–8. Available from: http://dx.doi.org/10.2136/sssaj1978.03615995004200030009x

27. Nkonge C, Balance GM. A sensitive colorimetric procedure for nitrogen determination in micro-Kjeldahl digests. J Agric Food Chem. 1982;30(3):416–20. Available from: http://dx.doi.org/10.1021/jf00111a002

28. Ereful NC, Jones H, Fradgley N, Boyd L, Cherie HA, Milner MJ. Nutritional and genetic variation in a core set of Ethiopian Tef (*Eragrostis tef*) varieties. BMC Plant Biol. 2022;22(1):220. Available from: http://dx.doi.org/10.1186/s12870-022-03595-9

29. Biswas AK, Sahoo J, Chatli MK. A simple UV-Vis spectrophotometric method for determination of β-carotene content in raw carrot, sweet potato and supplemented chicken meat nuggets. LWT Food Sci and Tech. 2011; 44:1809–13.

30. R Core Team. R: A language and environment for statistical computing. R Foundation for Statistical Computing, Vienna, Austria. 2021. Available from: https://www.R-project.org/

31. Wickham, H. ggplot2: Elegant Graphics for Data Analysis. Springer-Verlag New York, 2016.

32. Inglis PW, Pappas CR, Resende LV, Grattapaglia D. Fast and inexpensive protocols for consistent extraction of high quality DNA and RNA from challenging plant and fungal samples for high-throughput SNP genotyping and sequencing applications. PLoS One. 2018;13(10):e0206085. Available from: http://dx.doi.org/10.1371/journal.pone.0206085

33. Andrews, S. (2010). FastQC: A Quality Control Tool for High Throughput Sequence Data [Online]. Available from http://www.bioinformatics.babraham.ac.uk/projects/fastqc/

34. Cunningham F, Allen JE, Allen J, Alvarez-Jarreta J, Amode MR, Armean IM, et al. Ensembl 2022. Nucleic Acids Res. 2022;50(D1):D988–95. Available from: http://dx.doi.org/10.1093/nar/gkab1049

35. Li H, Durbin R. Fast and accurate short read alignment with Burrows-Wheeler transform. Bioinformatics. 2009;25(14):1754–60. Available from: http://dx.doi.org/10.1093/bioinformatics/btp324

36. Danecek P, Bonfield JK, Liddle J, Marshall J, Ohan V, Pollard MO, Whitwham A, Keane T, McCarthy SA, Davies RM, Li H. Twelve years of SAMtools and BCFtools. GigaScience. 2021; 10(2). Available from https://doi.org/10.1093/gigascience/giab008

37. Poplin R, Ruano-Rubio V, DePristo MA, Fennell TJ, Carneiro MO, Van der Auwera GA, et al. Scaling accurate genetic variant discovery to tens of thousands of samples. bioRxiv, 2017: 201178. DOI: 10.1101/201178

38. Danecek P, Auton A, Abecasis G, Albers CA, Banks E, Depristo MA, et al. Richard Durbin and 1000 Genomes Project Analysis Group. The Variant Call Format and VCFtools. Bioinformatics. 2011.

39. Cingolani P, Platts A, Wang LL, Coon M, Nguyen T, Wang L, et al. A program for annotating and predicting the effects of single nucleotide polymorphisms, SnpEff: SNPs in the genome of *Drosophila melanogaster* strain w1118; iso-2; iso-3. Fly (Austin). 2012;6(2):80–92. Available from: http://dx.doi.org/10.4161/fly.19695

40. Raj A, Stephens M, Pritchard JK. FastSTRUCTURE: Variational inference of population structure in large SNP data sets. Genetics. 2014;197(2):573–89. Available from: http://dx.doi.org/10.1534/genetics.114.164350

41. Cook DE, Andersen EC. VCF-kit: Assorted utilities for the variant call format. Bioinformatics. 2017;btx011. Available from: http://dx.doi.org/10.1093/bioinformatics/btx011

42. Wickham H, Averick M, Bryan J, Chang W, McGowan LD, François R, et al. “Welcome to the tidyverse.” Journal of Open Source Software, 2019; 4(43), 1686. doi:10.21105/joss.01686.

43. McWilliam H, Li W, Uludag M, Squizzato S, Park YM, Buso N, Cowley AP, Lopez R Analysis Tool Web Services from the EMBL-EBI. (2013). Nucleic acids research 2013 Jul;41(Web Server issue):W597–600 doi:10.1093/nar/gkt376

44. Howeler R. Sustainable soil and crop management of cassava. A reference manual. Centro Internacional de Agricultura Tropical CIAT Publication No. 389. Cali, Columbia: Centro Internacional de Agricultura Tropical. CIAT; 2014.

45. Hu W, Sarengaowa, Guan Y, Feng K. Biosynthesis of Phenolic Compounds and Antioxidant Activity in Fresh-Cut Fruits and Vegetables. Front Microbiol. 2022; 25(13):906069. doi: 10.3389/fmicb.2022.906069.

46. Anantharaju, PG, Gowda PC, Vimalambike, MG et al. An overview on the role of dietary phenolics for the treatment of cancers. Nutr J. 2016; 15, 99. https://doi.org/10.1186/s12937-016-0217-2

47. Grotz N, Fox T, Connolly E, Park W, Guerinot ML, Eide D. Identification of a family of zinc transporter genes from Arabidopsis that respond to zinc deficiency. Proc Natl Acad Sci U S A. 1998;95(12):7220–4. Available from: http://dx.doi.org/10.1073/pnas.95.12.7220

48. Chen WR, Feng Y, Chao YE. Genomic analysis and expression pattern of OsZIP1, OsZIP3, and OsZIP4 in two rice (*Oryza sativa* L.) genotypes with different zinc efficiency. Russ J Plant Physiol. 2008;55(3):400–9. Available from: http://dx.doi.org/10.1134/s1021443708030175

49. Food Data Central. United States Department of Agriculture. Available online: https://fdc.nal.usda.gov/fdc-app.html#/food-details/168875/nutrients. Accessed on Jan 15, 2023

50. Cakmak I, Pfeiffer WH, McClafferty B. REVIEW: Biofortification of durum wheat with zinc and iron. Cereal Chem. 2010;87(1):10–20. Available from: http://dx.doi.org/10.1094/cchem-87-1-0010

51. Das P, Adak S, Lahiri Majumder A. Genetic manipulation for improved nutritional quality in rice. Front Genet. 2020;11:776. Available from: http://dx.doi.org/10.3389/fgene.2020.00776

52. Swamy BPM, Samia M, Boncodin R, Marundan S, Rebong DB, Ordonio RL, et al. Compositional analysis of genetically engineered GR2E “Golden Rice” in comparison to that of conventional rice. J Agric Food Chem. 2019;67(28):7986–94. Available from: http://dx.doi.org/10.1021/acs.jafc.9b01524

53. Palanog AD, Calayugan MIC, Descalsota-Empleo GI, Amparado A, Inabangan-Asilo MA, Arocena EC, et al. Zinc and iron nutrition status in the Philippines population and local soils. Front Nutr. 2019;6:81. Available from: http://dx.doi.org/10.3389/fnut.2019.00081

54. Mulder EG. Nitrogen – Magnesium Relationships in Crop Plants. Plant and Soil. 1956;7(4):34176.

55. Paradisone V, Navarro-León E, Ruiz JM, Esposito S, Blasco B. Calcium silicate ameliorates zinc deficiency and toxicity symptoms in barley plants through improvements in nitrogen metabolism and photosynthesis. Acta Physiol Plant. 2021;43(12). Available from: http://dx.doi.org/10.1007/s11738-021-03325-y

56. Paine JA, Shipton CA, Chaggar S, Howells RM, Kennedy MJ, Vernon G, et al. Improving the nutritional value of Golden Rice through increased pro-vitamin A content. Nat Biotechnol. 2005;23(4):482–7. Available from: http://dx.doi.org/10.1038/nbt1082

57. Swamy BPM, Senior Scientist I – Rice Breeding and Biofortification. Personal Communication. International Rice Research Institute. 2022.

58. Zhang B, Liu J, Cheng L, Zhang Y, Hou S, Sun Z, et al. Carotenoid composition and expression of biosynthetic genes in yellow and white foxtail millet [*Setaria italica* (L.) Beauv]. J Cereal Sci. 2019;85:84–90. Available from: http://dx.doi.org/10.1016/j.jcs.2018.11.005

59. Brotman Y, Llorente-Wiegand C, Oyong G, et al. 2021. The genetics underlying metabolic signatures in a brown rice diversity panel and their vital role in human nutrition. Plant J. 106(2):507–525. doi: 10.1111/tpj.15182.

60. Moore, R., Jackson, K., & Minihane, A. (2009). Green tea (*Camellia sinensis*) catechins and vascular function. British Journal of Nutrition, 102(12), 1790–1802. doi:10.1017/S0007114509991218

61. Srinivasulu C, Ramgopal M, Ramanjaneyulu G, Anuradha CM, Suresh Kumar C. Syringic acid (SA) _ A review of its occurrence, biosynthesis, pharmacological and industrial importance. Biomed Pharmacother. 2018; 108:547–557. Available from: http://dx.doi.org/10.1016/j.biopha.2018.09.069

62. Imran, M.; Salehi, B.; Sharifi-Rad, J.;, et al. Kaempferol: A Key Emphasis to Its Anticancer Potential. Molecules 2019, 24, 2277. https://doi.org/10.3390/molecules24122277

63. Wittkopp PJ, Haerum BK, Clark AG. Evolutionary changes in cis and trans gene regulation. Nature. 2004;430(6995):85–8. Available from: http://dx.doi.org/10.1038/nature02698

64. Ding Y, Zhu J, Zhao D, Liu Q, Yang Q, Zhang T. Targeting Cis-regulatory elements for rice grain quality improvement. Front Plant Sci. 2021;12:705834. Available from: http://dx.doi.org/10.3389/fpls.2021.705834

65. Kakei Y, Masuda H, Nishizawa NK, Hattori H, Aung MS. Elucidation of novel cis-regulatory elements and promoter structures involved in iron excess response mechanisms in rice using a bioinformatics approach. Front Plant Sci. 2021;12. Available from: http://dx.doi.org/10.3389/fpls.2021.660303

66. Tirosh I, Reikhav S, Levy AA, Barkai N. A yeast hybrid provides insight into the evolution of gene expression regulation. Science. 2009;324(5927):659–62. Available from: http://dx.doi.org/10.1126/science.1169766

67. Ereful NC, Laurena A, Liu L-Y, Kao SM, Tsai E, Greenland A, et al. Unraveling regulatory divergence, heterotic malleability, and allelic imbalance switching in rice due to drought stress. Sci Rep. 2021;11(1). Available from: http://dx.doi.org/10.1038/s41598-021-92938-x

